# KERA: Analysis Tool for Multi-Process, Multi-State Single-Molecule Data

**DOI:** 10.1101/2021.01.04.425319

**Authors:** Joseph Tibbs, Mohamed Ghoneim, Colleen C. Caldwell, Troy Buzynski, Wayne Bowie, Elizabeth M. Boehm, M. Todd Washington, S. M. Ali Tabei, Maria Spies

## Abstract

Molecular machines within cells dynamically assemble, disassemble, and reorganize. Molecular interactions between their components can be observed at the single-molecule level and quantified using colocalization single-molecule spectroscopy (CoSMoS), in which individual labeled molecules are seen transiently associating with a surface-tethered partner, or other total internal reflection fluorescence microscopy (TIRFM) approaches in which the interactions elicit changes in fluorescence in the labeled surface-tethered partner. When multiple interacting partners can form ternary, quaternary and higher order complexes, the types of spatial and temporal organization of these complexes can be deduced from the order of appearance and reorganization of the components. Time evolution of complex architectures can be followed by changes in the fluorescence behavior in multiple channels. Here, we describe the kinetic event resolving algorithm (KERA), a software tool for organizing and sorting the discretized fluorescent trajectories from a range of single-molecule experiments. KERA organizes the data in groups by transition patterns, and displays exhaustive dwell-time data for each interaction sequence. Enumerating and quantifying sequences of molecular interactions provides important information regarding the underlying mechanism of the assembly, dynamics and architecture of the macromolecular complexes. We demonstrate KERA’s utility by analyzing conformational dynamics of two DNA binding proteins: RPA and XPD helicase.

## INTRODUCTION

Molecular machines that replicate, repair and survey DNA for damage are often dynamic assemblies of macromolecules that form, disassemble, undergo modifications and exchange components. In fact, the bustle of activity around moving or stalled DNA replication forks is instrumental to the fidelity and robustness of DNA synthesis (see (1–3) for recent comprehensive reviews). Stretches of ssDNA bound by the Replication protein (RPA), the main ssDNA binding protein in eukaryotic cells, or its bacterial counterpart SSB are channeled to DNA replication, repair, recombination or signaling events (4,5). DNA damage bypass, nucleotide excision repair (NER), and base excision repair (BER) depend on the timely and coordinated arrival of enzymes either to allow stalled replication forks to synthesize DNA across from DNA lesions in the template strand during replication or to remove DNA lesions from the genome prior to replication (6–9). RNA processing during slicing, transcription and ribosome assembly and function also involves complex and dynamic nucleoprotein transactions (10,11). Within these multicomponent architectures, transient interactions take place, while complexes are formed, dissolved and rearranged. Quantifying and enumerating these interactions can be assisted by single-molecule analyses.

Colocalization single-molecule spectroscopy (CoSMoS) (12,13) refers to a class of techniques, which observe the dynamics of a molecule or complex within a diffraction-limited area. Commonly, fluorescence is used as a reporter to interrogate one or more aspects of the system, with the ability of multi-channel setups to observe distinct colors of fluorescence independently and simultaneously for the same location. In some cases, a fluorescent signal at a location implies the binding of a labeled molecule. If an immobilized binding partner is tethered to the surface, the kinetics of the binding interaction can be quantified by measuring the duration of the fluorescent events (14,15); this situation is shown schematically by **Figure 1a**. However, fluorescent probes can also be used to access structural information, such as conformational states (16–18). This is because fluorophores can vary in brightness due to protein induced fluorescence enhancement (PIFE) and quenching (PIFQ) (19,20), iron-mediated fluorescence quenching (21,22), Forster resonance energy transfer (FRET) between donor and acceptor fluorophores (23,24) and other mechanisms. **Figure 1b** shows a cartoon of a fluorophore-labeled protein exploring three distinct conformational states, resulting in discrete levels of fluorescence. However, the greatest strength of these techniques is the simultaneous measurement of fluorescence from two (25) or more (26) spectrally-distinct channels. In these studies, the interaction of more than one labeled species (whether by binding, conformational change, or other fluorescent reporting) can be correlated in real time. In the study of PCNA toolbelt formation (27), for example, two labeled species were colocalized to produce trajectories like what is illustrated by **Figure 1c**, but three-channel TIRF is also employed, such as in a study which investigated spliceosome complex formation (28).

**Figure 1.**
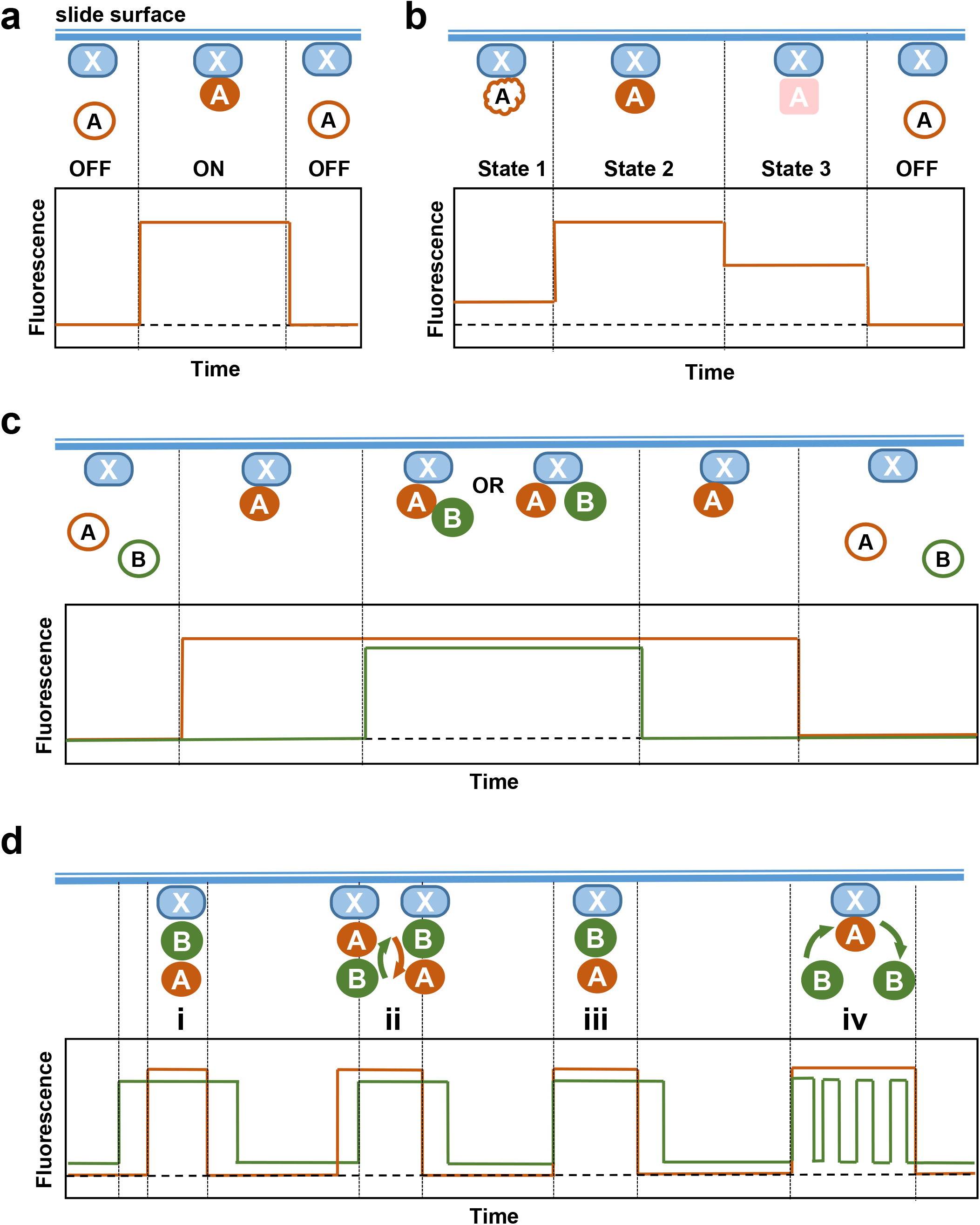
Schematics of possible evolution of single-molecule fluorescence over time for typical model systems. **(a)** A fluorescently-labeled species binds to an unlabeled immobilized partner and causes an increase in fluorescence at that location for the duration of the binding event. Measurement of the dwell times in ON and OFF state over a large number of binding events informs on the dissociation and association rate, respectively. **(b)** A macromolecule labeled with an environmentally sensitive fluorophore transitions between conformational states, which are observed as three distinct levels of fluorescence, low (state 1), high (state 2) and intermediate (state 3), respectively. **(c)** A ternary complex forms, identified through multi-channel imaging of species labeled with spectrally-distinct fluorophores. Conclusions regarding the architecture of the complex may be derived from the binding/dissociation sequence and an a priori knowledge of the interaction sites on the macromolecules X, A and B. From a single fluorescent event alone, one may conclude that either A bridges the interaction between X and B, or A and B bind to X simultaneously and independently. **(d)** Two-channel experiments can report on the architecture of the ternary complex involving a surface tethered molecule and its two interacting partners labeled with fluorescently distinct fluorophores. Similar to panel (c), a knowledge of the interaction sites on X, A and B are required to make an unambiguous determination of possible architectures. The schematics here assumes that the A and B interact with one another and share a common binding site on X.

Although **Figure 1a-c** presents the events as simple stepwise changes in fluorescence, in reality “steps” in the single-molecule trajectories can often be obscured by the noise and the closely spaced states may be difficult to distinguish. Although for binary ON-OFF time trajectories a simple thresholding procedure might allow the researcher to distinguish the states, some signals (as in **Figure 1b**) require more complicated models. Experimental trajectories are idealized whereby each segment of a trajectory is assigned to one of a few discrete states. Multiple analysis software packages have been developed to extract the state information from single-molecule trajectories (29–37). These programs apply methods including analysis of the noise, Hidden Markov Modeling (HMM) and Bayesian statistical approaches to translate each trajectory in an ensemble into a time series of discrete states (37).

Idealized trajectories contain important structure-functional information. At their most general, these models show the patterns of binding and dissociation of the tagged molecules, or a macromolecular complex exploring its conformational space. In a single-channel setup, intermediate states may indicate states of specific conformations or architectures (**Figure 1b**). A similar type of fluorescence pattern may be observed when multiple fluorescent molecules can bind to and dissociate from a surface-tethered partner. In a multi-channel setup, the binding molecules may bind in complexes or individually (**Figure 1c&d**). There may be a preferred order or a tendency for one molecule’s binding to inhibit binding with the second partner (27,38). As **Figure 1c** demonstrates, even a simple two-channel binding experiment can reveal important information about the system when the order of binding is considered (38). Without explicit prior knowledge of the patterns to look for, classifying the events into meaningful categories can be difficult. While various step finder programs help to determine which binding states are present in the trajectories, the binding behavior implied by the transitions between states are harder to identify. In addition, the length of time spent at each state, the dwell time, is a statistical quantity which can be used to determine important information about the kinetics of the binding (14). In multi-channel studies, where the duration of each state may depend on the state of other binding partners, there is as yet no simple way of organizing the data to discriminate between the possible multi-color transitions. Finally, correlating the information contained in two or more discretized data channels allows for a more complete picture of the interactions than an analysis of any channel individually.

Here, we present the kinetic event resolving algorithm (KERA), designed to analyze multi-channel and multi-state data. The purpose of this program is to search large datasets for patterns of transitions between discrete states, and to allow the researcher to study events of these patterns in greater detail, to determine if they are the artifacts of chance or the result of biochemical preference. In some cases, this allows properties of rarer events to be identified, beyond the broad ensemble averages. Thus, the information that the trajectory analysis provides is beyond mere state identification. It identifies the time at which each transition occurs, and how long the system remains in each state before transitioning. It analyzes the order and sequence of events in multi-channel and multi-process data, highlighting possible binding mechanisms and architectures of macromolecular complexes. KERA has been developed to automatically read the discretized time series data and identify and classify all the transitions and their correlations. In addition, the user may define custom transition patterns using a simple user interface or the power of regular expressions, giving KERA a versatility which extends to a wide variety of systems. Although KERA was designed with biochemical systems in mind, it has the ability to analyze discretized time data from any source which is formatted correctly. The scripts for KERA analyses are written in MATLAB and are available through a public GitHub repository at https://github.com/MSpiesLab/KERA. Ultimately, KERA aims to be an open-source, user-friendly MATLAB suite, which simplifies identification of discrete-state patterns and extraction of dwell times in complex multi-channel data.

## METHODS

### MATLAB Suite

KERA was designed as a suite of MATLAB code (version 2020a) which is available on GitHub at https://github.com/MSpiesLab/KERA, along with full documentation. The code consists of several main scripts, which in turn call a variety of functions included in a sub-folder. To run KERA, only the “openKERA.m” file is needed on the front end. This creates a window which utilizes MATLAB’s built-in GUI creation tools, such as buttons, menu options, and graphics. Some of the scripts create temporary new GUI windows which are used to examine the traces or alter the discretization before being closed. The workflow starts with importation, followed by optional inspection of the traces and discretization, before creating an analysis profile for the data. A battery of default search patterns are included for ease of use, but after they are complete the user can define custom searches in a variety of ways, to be appended to the current results. If new data are imported, they can be appended to the existing data, and all current search results updated to include the new data. Search parameters can be copied to new datasets and run there, as long as the datasets have compatible numbers of channels and states. After analysis, dwell-time data are available for export to csv files, while the MATLAB arrays held within the output structure can also be copied and pasted into a spreadsheet application. Finally, the GUI window itself and all the data it contains is easily exported so that an analysis session and all changes made to the data can be recovered at a later time or in another KERA session.

### Analysis of XPD conformational dynamics during interactions with damaged DNA substrates

Intensity-time trajectories were selected from experiments, which used Cy3-labeled XPD and Cy5-labeled DNA, as previously reported by Ghoneim and Spies (22). For KERA analysis, the data used came from the trials in which the Cy5-labeled DNA substrate contained a cyclobutane pyrimdine dimer (CPD) and biotinylated, Cy3-labeled XPD helicase was tethered to the surface. Changes in the Cy3 fluorescence were reporting on the conformational transitions that bring the Cy3-labeled ARCH domain of XPD close to and away from the FeS domain, while appearance and disappearance of the Cy5 signal signified DNA binding and dissociation, respectively. After extracting intensity-time trajectories from raw videos, a set of selection rules was applied to decide on the trajectories that would be used in the analysis. Those chosen had an average intensity profile which was stable over time, without systematic fluorescence level drift of constant states. For Cy3 (ARCH domain) trajectories, at least two conformational transitions had to be present in the trajectory. Trajectories with signal-to-noise ratio of less than 5:1 were excluded. All accepted trajectories ended with irreversible single-step photobleaching from a higher fluorescence state, and were longer than 10 seconds before trimming. Previously, a custom MATLAB user interface was used for normalization, baseline correction and 50% thresholding to separate the open and closed states of XPD ARCH domain (22). Here, traces were normalized and baseline-corrected by using a custom MATLAB script (normalizeTrajectorySet) to select regions of baseline and the highest fluorescent state which lasted longer than 3 frames. The trajectories were also trimmed to exclude regions after Cy3 photobleaching. hFRET (32), developed by the Gonzalez lab for use in kinetically heterogeneous systems, was the chosen discretization software. The hFRET method requires that all traces be the same length upon import; the traces were thus padded at the end with 0 values to a uniform length. hFRET was run using an Hidden Markov Model with one dimension, 1 sub-population per state, shared emissions, priors manually set, and analyzing the absolute amplitudes. For the Cy3 data, four states were used in the model, with the first being assigned to the 0 padding; the priors were set to [0, 0.2, 0.5, 0.9]. For the Cy5 data, three states were used, at [0, 0.2, 0.9]. After the analysis was complete, the discretized Viterbi path trajectories were imported to KERA. The padding state was removed from all trajectories (or assigned to the baseline, state 1, in the few occasions it occurred in the middle of the data), resulting in a 3-state model for the Cy3 signal reporting on the XPD conformations and 2-state model for the Cy5 reporting on the DNA binding.

### Analysis of the RPA conformational dynamics

Fluorescence intensity trajectories of MB543-labeled RPA molecules on immobilized ssDNA were extracted from a subset of the raw microscope movies previously described in Pokhrel, Caldwell et al. (12). The trajectories selected for analysis were those which had no fluorescence in the first 30 seconds (before fluorescent proteins were added), contained at least two state transitions, and had a raw signal-to-noise ratio of at least 4. The trajectories were shifted using a baseline-thresholding method to align their lowest state and then normalized by their highest state. The trajectories were trimmed and split to analyze each portion of the experiment separately: one split from frames 300-1,200 of acquisition (after RPA addition) and one from frames 1,200 to 2,100 (after RAD52 addition or buffer wash). These segments were then paired with a synthetic “donor” segment which would result in a calculated FRET value which was equal to the original segment. As a result, when these pairs were imported to ebFRET for analysis, the FRET signal analyzed was equal to the normalized trace. ebFRET was used to obtain a four-state model fitted to each ensemble of traces which were then exported to the single-molecule dataset (smd) format. The discretized Viterbi path time series were imported to KERA, which was used to extract the dwell times of all non-edge states throughout the ensemble. Histograms were plotted and analyzed to extract rate constants using GraphPad Prism 7 as described in detail in (39).

## RESULTS

### Overview of the KERA system

We have developed KERA, a MATLAB suite, united by a simple GUI window which allows the user to run analyses and export the results. The first step in the analysis is to import the data and convert it to a unified format. **Figure 2a** shows an example of individual events selected from noisy single-molecule data contained in pairs of colocalized fluorescence trajectories (intensity time series). Before they are imported to KERA, the trajectories must be turned into time series of discrete states, a process referred to as idealization (**Figure 2b**). Idealization can be carried out using any appropriate state-finding package. Below, we show two examples one utilizing hFRET (32) and the second utilizing ebFRET (31). After importing the idealized trajectories, KERA’s main functionalities are the ability to sort events and transitions by categories and allow the researcher to search the set for transitions and conformations of interest. **Figure 2c-e** provides an example of the type of information this categorization provides. If this were a binding experiment of two distinctly-labeled species (Red and Green), the prevalence of events in which Red or Green bound first might imply a preferred binding order or architecture. For each such category of transitions specified, dwell time data can be visualized using the built-in plotting tools, and basic fitting methods allow for a fast estimate of kinetic information in each group of transitions (**Figure 2e**). KERA’s other features include a data browser to view and interact with the discretized and raw data, as well as a dwell time summary which organizes each channel’s state dwell times into easily-accessible data tables.

**Figure 2:**
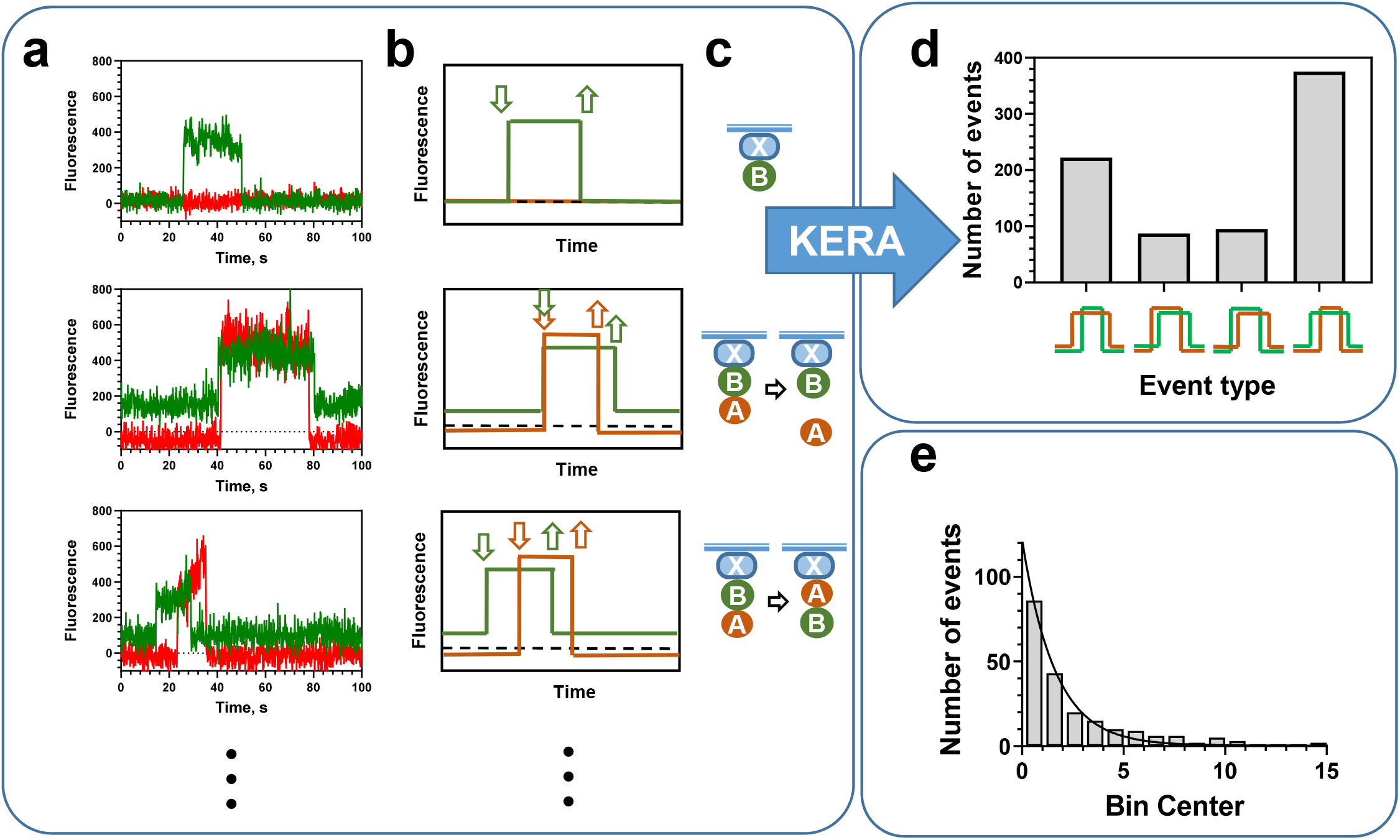
Overview of the KERA workflow and features. **(a)** Examples of events from representative raw intensity trajectories from the data described in (27). **(b)** The same events shown as idealized trajectories. **(c)** Interpretation of the respective molecular architectures. Top: a binary complex formation between surface tethered molecule X and fluorescently-labeled molecule B. Middle: a binary complex between fluorescently-labeled molecules A and B arrives to the surface-tethered X; A dissociates before B suggesting that in the trimeric complex B bridges the interaction between X and A. This scenario assumes that A and B interact with the same site on X. Bottom: the arrival of B followed by A results in a ternary complex. However, in this scenario, there is only one binding site on X, so the departure of B first implies a reversal of the binding partner, as found in the PCNA Toolbelt-Rev1 Bridge switch observed in (27). Other interpretations, dependent on the system, are possible. **(d)** The idealized data are imported to KERA. An example analysis compares the prevalence of certain transition patterns (event classifications) within the data (the numbers are from (27)). **(e)** For a given event type, the dwell times are plotted as a histogram and an exponential fit is applied to extract estimated kinetic data.

### KERA Data Workflow

In single-molecule studies, many systems of interest may be interrogated with the use of fluorescent probes such that the intensity of the signal is correlated with some useful property of the system (binding or conformational change, for example). Single-molecule Total Internal Reflection Microscopy (smTIRFM) is well-suited to this task, and many papers describing methods to detect and quantify colocalized fluorescence are available (26,40–42). For the simple binary models, such as when molecules are either bound or unbound with no intervening states, it is common to use a thresholding operation to define the regions of “ON” fluorescence and “OFF” fluorescence. For other models, software such as ebFRET (31,43), QuB (44), HaMMY (29), hFRET (32), and SMACKS (45) are powerful discretization tools. Although these methods (with the exception of QuB) were developed specifically for FRET signals, they can be adapted to analyze fluorescence trajectories from single-color experiments and experiments with independent fluorescence signals. These are not the only methods of discretizing a fluorescence time series, but KERA contains a user interface for importing datasets created by QuB, ebFRET, hFRET and HaMMY software packages (**Figure 3a-c**). In addition, any data which exist as a time series of integer states may be formatted to match KERA’s raw input format, which is described thoroughly by the documentation included with the code, and instructions are also included for altering the source code to allow for new import formats.

**Figure 3:**
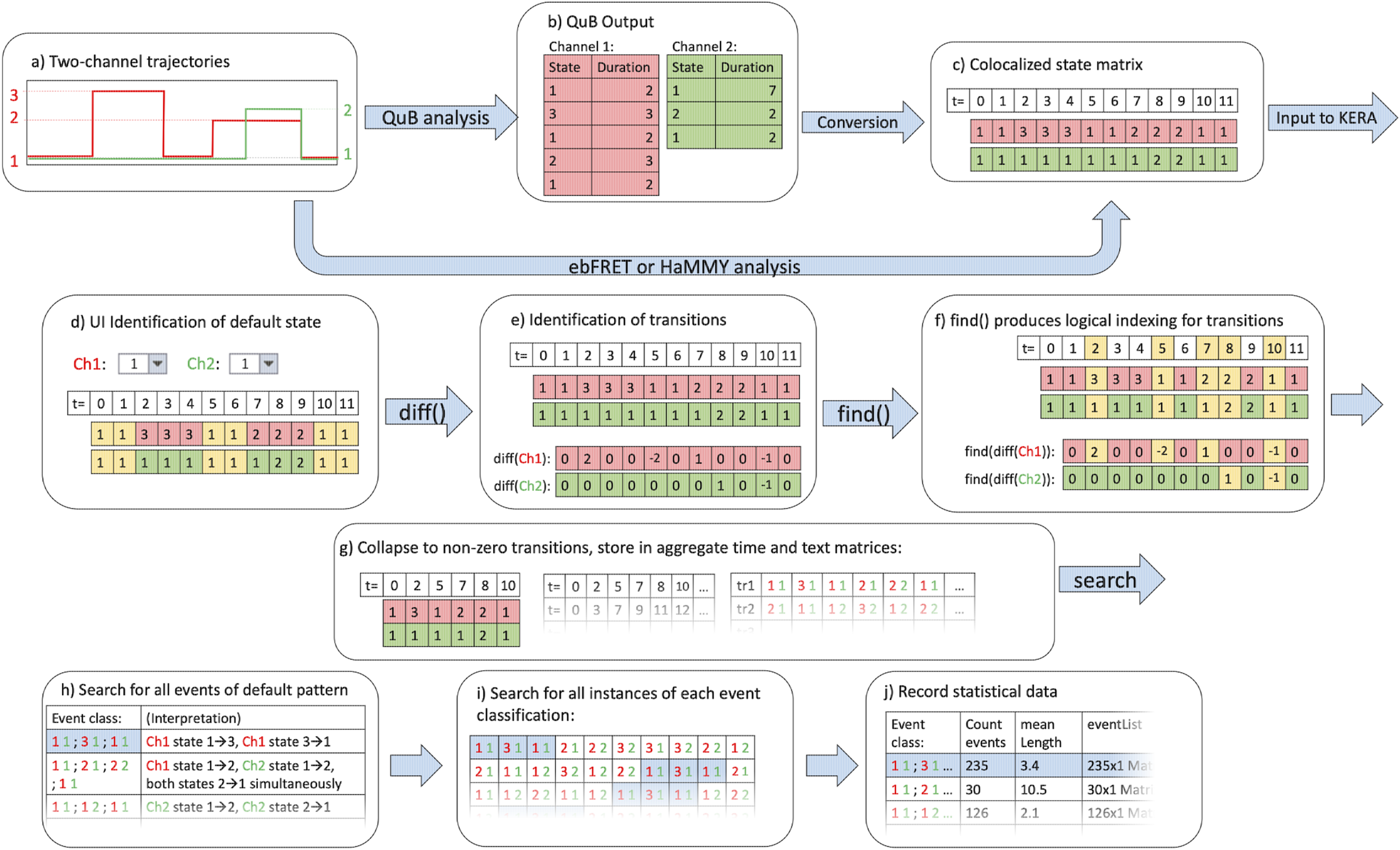
Flow of the program structure. A flow chart is showing the program’s default search method on a single two-channel multistate trajectory: **(a)** A cartoon trajectory with two channels is idealized, either in QuB or ebFRET. The red channel has three states and the green channel has two states. Although only three trajectories are shown, hundreds or thousands would be used in a given analysis. **(b)** QuB analysis would produce two tables listing the states and durations for each channel in the trajectory; this is converted by the program into **(c)** The output format of ebFRET, and the common input format for KERA **(d)** The user identifies the “default state”, which KERA will search for as marking the beginning and ending of kinetic events in the initial search. Events not matching this default state can later be specified by custom search. **(e)** The diff() built-in function locates all instances of a change in state. **(f)** These indices are used to identify time points of interest; all other entries are redundant **(g)** The collapsed matrix is shown on the left. The middle matrix is the aggregate timestamp matrix, where time data is stored for all trajectories. The right matrix is the aggregate state matrix, where state data is stored as text strings delimited by semicolons. **(h)** The condensed state matrix is searched to find all unique event classifications, which are recorded in a list. For each of these, KERA will (i) search for events matching that classification, and (j) record ensemble and individual event data about those events in a table.

Once the data are imported, the analysis organizes them into a searchable matrix and pairs colocalized data from disparate files into a single trajectory. **Figure 3** shows this import process for a simple, cartoon trajectory pair (**Figure 3a**). The molecule in the red channel has three binding states (off, semi-bound, and bound) while the green channel has only an on- and off-state. Colocalized trajectories, or the trajectories describing data from the same location but in different fluorescence channels, are organized as successive rows in a matrix. Because most events of interest occur between instances of the baseline or default state of the system, the user has the option to specify which state each channel exists in when there is no activity (**Figure 3d**); this allows the program to highlight all departures from that state in the later analysis. MATLAB’s built-in diff() function is used on each trajectory to identify all changes in state with a non-zero entry; all time points in which the state was constant are replaced by 0 entries (**Figure 3e**). Using the find() (location of non-zero elements) function allows these vectors to be collapsed to a minimal form, where only transitions between binding states and the timestamp of each transition are retained (**Figure 3f**). Each of these minimal vectors is then added as an entry in the aggregated transition or timestamp cell arrays, allowing for quick reference between transitions and the time at which they occurred (**Figure 3g**).

Each trajectory is now encoded by an array of integers corresponding to the state of each channel. The program then scans the dataset for matches to the user-defined default binding behavior and makes a list of all unique event classifications, accomplished by applying MATLAB’s unique() function to the results of the initial search. In essence, a pattern which begins with the default state and returns to it after any number of transitions is searched for in the data. This default state may be set by the user, but by default it is assumed that non-fluorescence in all states is the ‘fully unbound’ or ‘default’ state to which the system returns eventually. All matches are recorded in a list, and all unique elements of that list are recorded as separate classifications. A classification is defined as a specific ordered set of transitions forming an event; a few possible such classifications are shown in **Figure 3h**. As demonstrated in **Figure 2**, event classifications depend on the binding or transition order. Each of these classifications becomes the subject of a subsequent detailed search of the data, and it is the classifications that direct the organization of the final output of the program. In addition to these automatically-defined classifications, the user has the ability to define custom binding patterns to search for. The user interface for this allows for a limited system of wildcard and token states, increasing the versatility of these user-defined patterns. For more complex patterns, full support of regular expressions allows the user to define search strings using raw regex input, a powerful tool for pattern recognition.

Once an event classification is defined, every event matching the description of that classification is catalogued and analyzed by the code in a single, comprehensive table. This means that, for a given classification, the program searches the transition matrix for events matching that description (**Figure 3i**). By cross-referencing the indices of search results with the time data matrix, the timestamps of all events are recovered efficiently. This produces time data about the events individually and as an ensemble within that classification, which are recorded in a table. Each classification’s ensemble data receives its own row in that table, for convenient comparison (**Figure 3j**). Events which are not completed before the end of the trace, or which begin before the start of the trace, are ignored, since reliable time data cannot be attached to that event.

### Visualization and Post-Processing

Although the program identifies all unique event classifications through this method, it is often more of interest to examine partial events, or specific single transitions, regardless of where they may occur in the set. For this purpose, KERA will also catalogue all instances of each configuration (multi-channel combination of states) and each single-channel transition type, along with their dwell time distributions. A custom search feature allows the user to specify a string of states in a channel, and thus filter the configurations by the following or preceding state. This is a useful in cases where the kinetics of a given state may exhibit hysteresis: the state being transitioned to (or from) impacts the dwell time of the state of interest.

The resulting sets of dwell times may be analyzed to determine important kinetic information about the system. The rate constants for binding and dissociation may be determined from histograms of the dwell times obtained by this analysis (38,46). Thorough treatment of the kinetics of binary binding possibilities have been reported in numerous previous studies. However, correlating the kinetic and thermodynamic data in separate channels (such as transitions between conformational states dependent on the bound and unbound states of a separate molecular component) is a more difficult task. This is, in part, because of the difficulty in obtaining dwell times which are specific to transitions between particular multi-channel system states. KERA, with its ability to categorize transitions and states and thus differentiate between different binding behavior in multi-channel experiments, collects dwell times into groups which are more amenable to this kind of analysis. For example, in the study of the interactions of XPD protein with damaged DNA substrates, the differential kinetics of DNA binding were correlated with the conformational state of a domain on the protein (see XPD case study in Results). In the study of the ternary complexes formed by PCNA, Polη, and Rev1, the dwell times associated specifically with each architecture could be extracted with KERA, taking into account information from both channels to distinguish system states (27).

For the dwell times of any given event classification, the program can optionally display basic dwell time histograms and cumulative distributions, to aid in kinetics evaluation. This creates a graph similar to those shown in **Figure 4**. For both the histograms and the cumulative distributions, two options are available: the dwell times may be binned linearly, to form an exponential distribution, or they may be binned logarithmically. In the latter case, the data points are taken to be the natural logarithm of each dwell time. The horizontal scale changes to reflect this. The advantage of this approach is that it becomes more visually apparent whether a single-or a double-exponential model is most suited to the data (47). This interface is illustrated by **Figure 4**. In addition to histograms, the program can also display and fit models to cumulative distributions, which are more useful in cases where fewer data points are available for the creation of a robust histogram.

**Figure 4:**
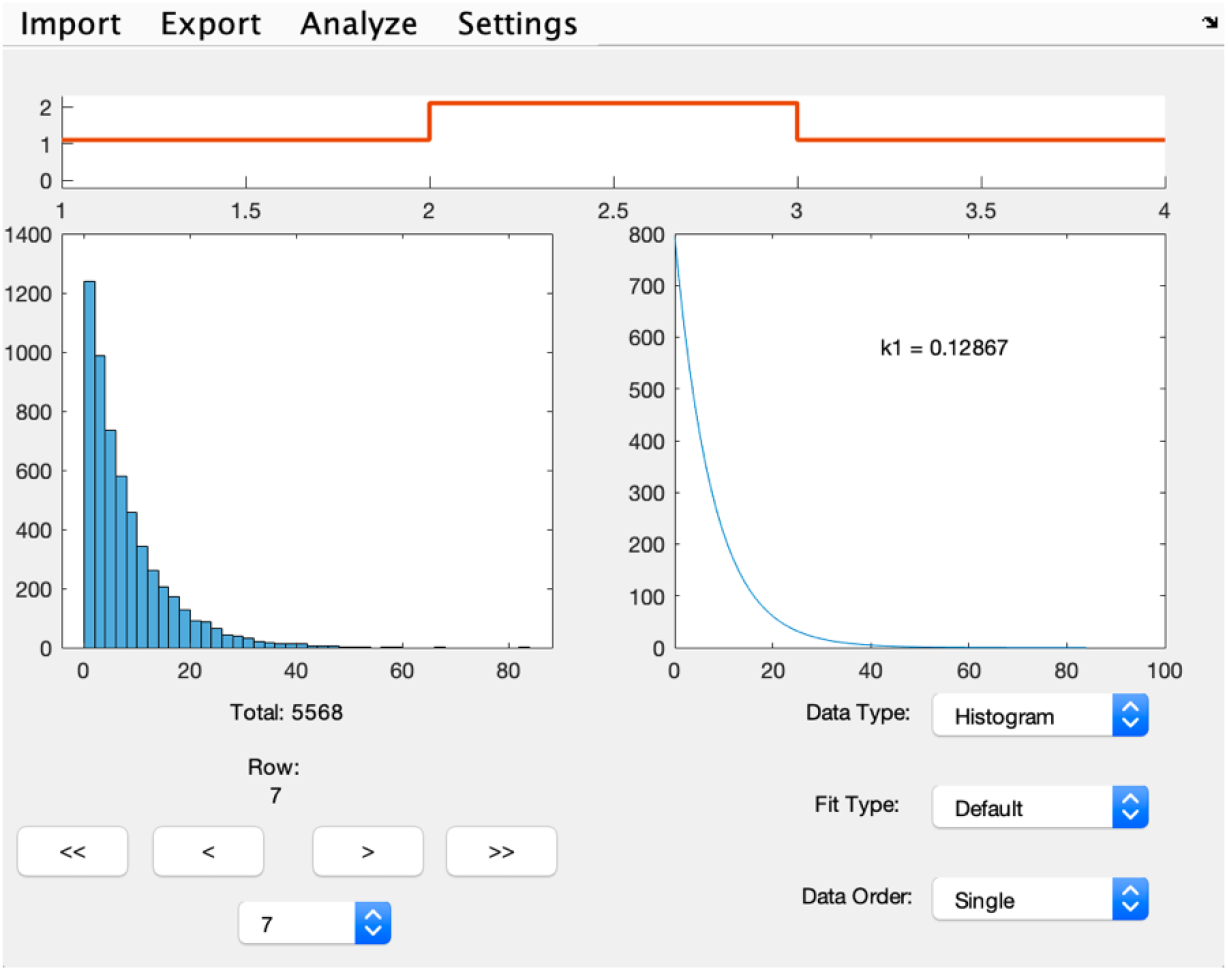
Program Output screen. An illustration of the graphical output of KERA, which displays select data from the larger set of results. For each event classification, a cartoon depiction of a representative transition is shown at the top of the window (here, channel 2 changes from state 1 to state 2 and back). The total number of such events is counted and displayed below a histogram of the dwell times of the event. The arrow buttons allow the user to move between event classifications. The fitting options on the right change the way the histogram is displayed and fitted. This analysis was run on simulated two-state two-channel data.

Examining these histograms and the outputted best-fit curve data shows at a glance which processes are kinetically faster, and which ones may depend on more than one rate constant. This portion of the suite is not meant to rigorously determine the kinetics of the interactions occurring in solution; it is more a tool for the researcher to examine their data without leaving the MATLAB environment in which the dwell time data is calculated. As noted, all data are available in a format which may be copied directly to a spreadsheet, allowing statistical analyses by dedicated data fitting packages.

Ultimately, these capabilities of searching data for event classifications and then extracting dwell time data from those events are meant to aid the researcher in discovering biochemically-relevant binding behaviors, as well as extracting quantitative information. In a binding experiment with thousands of colocalized pairs, the interpretation of the ensemble’s behavior is made more complex by the vast quantity of data and binding events available. Even in a single-channel experiment, a model which has more than two possible states becomes onerous to interpret, because each dwell time may depend on the states which the system takes on before and after a given binding site. In addition, the states which are achieved may exhibit a preferred order, indicated in the data by a predominance of one binding pattern; without a method to catalogue and count all distinct patterns, locating such preferred patterns would be time-consuming and frustrating, even using the transition density plots which FRET data analysis packages provide. KERA allows many patterns to be discerned by a simultaneous scan through all of the data, and the ability to search through the data in a customizable way allows researchers to extract specific data about state-dependent dwell times. For researchers with experience in Regular Expressions, raw regex input allows for even greater flexibility in the patterns which are categorized. These advanced methods are described in the documentation, but KERA’s userfriendly interface allows for a default scan of a single-molecule dataset with minimal user input.

The two case studies included below focus on macromolecular interactions with a time evolution through discrete states modeled by a Markov process. Indeed, KERA provides tools and visualizations which make the analysis of Markovian processes simpler, but it is equally true that the core functionality of KERA does not assume any such processes are at work. Idealized data can come from any source, including ones which are non-Markovian or far from equilibrium, and KERA will still parse it to effect the classification and enumeration of the patterns contained in the data. Thus the identification of rare events as well as the discovery of events which occur more often than would be assumed in a Markovian system are both possible in KERA. Although examples of nucleoprotein interactions are most relevant to the scope of this paper and the original motivation of KERA, discrete-time state transitions from any system may be analyzed as long as the data is placed into MATLAB cell variables of the correct format.

### Case Study 1. Discerning the XPD helicase domain dynamics in free and DNA bound states

XPD (Xeroderma pigmentosum complementation group D) protein is a helicase which provides an important function in Nucleotide Excision Repair (NER) (48,49) due to its ability to verify DNA damage (50). In addition to superfamily 2 helicase motor domains (HD1 and HD2), and an iron-sulfur (FeS) containing domain, XPD contains a flexible ARCH domain, which protrudes from the HD1 creating a pore in the protein. DNA passes through this pore from the DNA binding site in the helicase core to the secondary DNA binding site located at the interface between HD1 and FeS domain (51-53). Using smTIRFM, Ghoneim and Spies (22) visualized motions of the ARCH domain. In this study, biotinylated XPD molecules site-specifically labeled with Cy3 dye at the ARCH domain were immobilized on the surface of a microscope slide. The presence of an endogenous FeS cluster in XPD caused distance-dependent Cy3 quenching when the ARCH domain was in “closed” conformations, resulting in discrete fluorescence changes. Direct correlation of ARCH domain motion and binding of DNA substrate was achieved by supplying the reaction with damage-containing DNA molecules labeled with Cy5, a dye spectrally distinct from the dye in the ARCH domain. Fluctuations in the Cy3 signal reported on the ARCH motion (i.e. opening and closing), while the abrupt increase and decrease in the Cy5 signal reflected DNA binding to and dissociation from surface-tethered XPD, respectively (see **Figure 5a** for a cartoon schematic of the experimental system and **Figure 5b** for representative two-color fluorescence trajectories).

**Figure 5:**
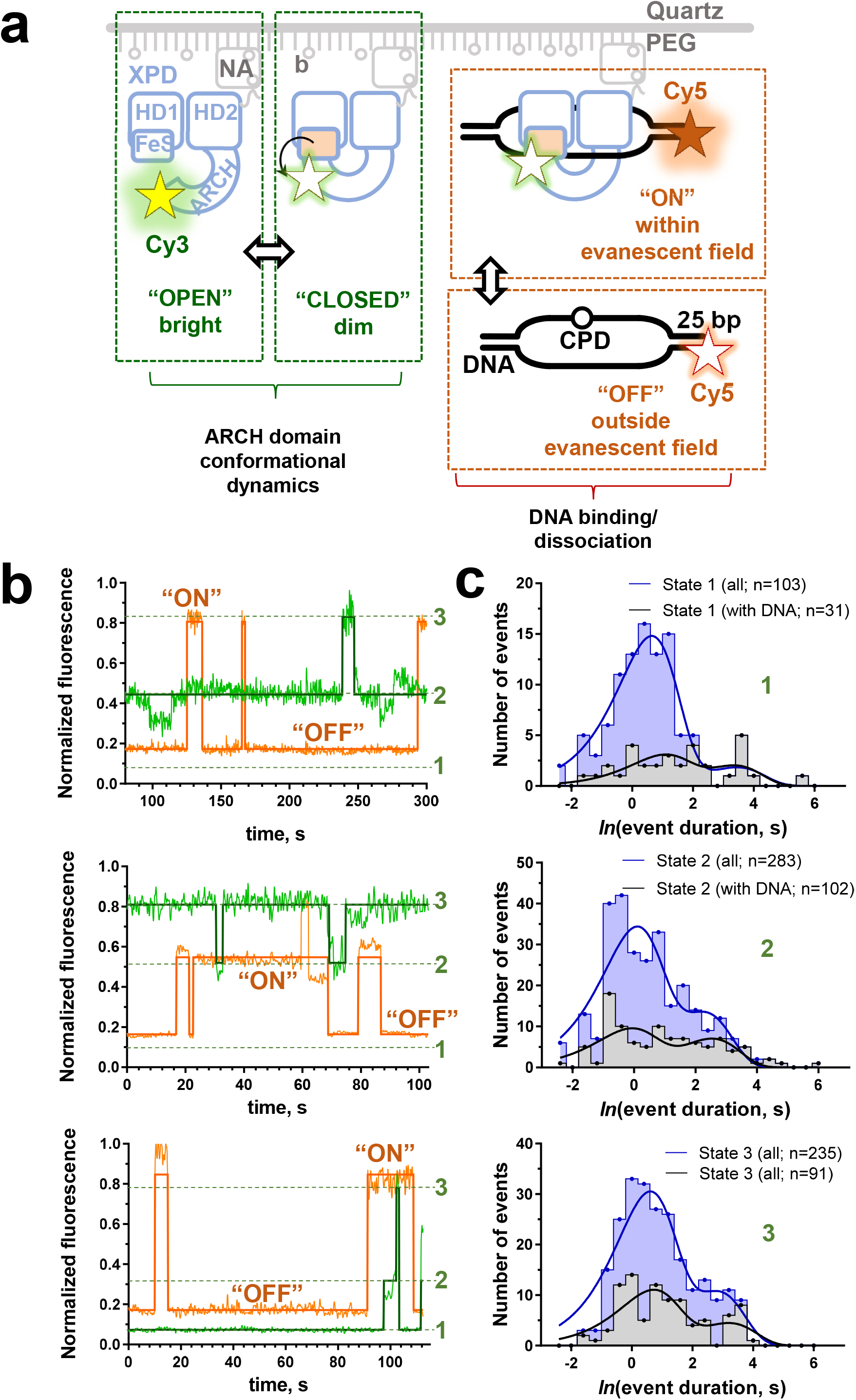
Representative discretized trace and molecular schematic from the XPD experiment. A subset of the original data from (22) was reanalyzed using KERA. **(a)** Experimental design. Biotinylated, Cy3-labeled XPD helicase is immobilized on the surface of the TIRFM flow cell. The states of the Cy3 fluorophore reflect the XPD ARCH domain conformations, as the proximity of the Cy3 dye site-specifically positioned in the ARCH domain to the endogenous ironsulfur cluster (FeS) results in the Cy3 quenching. The damaged DNA substrate was labeled with a Cy5 dye and only exhibited fluorescence when bound to the immobilized XPD at the slide surface. **(b)** Three sets of representative fluorescence intensity-time trajectories, each representing the fluorescence signal from a pair of colocalized Cy3 (green) and Cy5 (orange) fluorophores. The solid lines indicate the discrete states which the noisy trajectories were fit to using the hFRET. Two states “ON” (DNA bound to XPD) and “OFF” (free XPD) are observed in the Cy5 trajectories, while the Cy3 trajectory was fit with a three-state model (1 – “closed” ARCH, 2 – “intermediate/partially open” and 3 – “open” ARCH). **(c)** Dwell time distributions for each state of the ARCH domain. Natural logarithms of dwell times were binned in 0.4 intervals, plotted as histograms using GraphPad Prism software and fitted to double exponentials. For each state, blue distributions contain all events irrespective of whether DNA was present or absent, while grey distributions contain only a subset of dwell times when the DNA was present.

In the original study, the data in both Cy3 and Cy5 channels were analyzed using two state models, where “open” and “closed” states of the ARCH domain were assigned using 50% threshold, while the “bound” and “free” states of XPD corresponded to the presence and absence of the Cy5 signal. Based on the kinetics analysis, several states were identified within “open” and “closed” states. Here, we have re-analyzed a subset of the data that corresponded to the XPD interaction with damage-containing DNA. The data (N = 60 trajectories) were discretized using hFRET (see methods) and imported into KERA. A three-state model was chosen for the Cy3 conformational signal, and a two-state model was used for the Cy5 binding signal. The resulting analysis illustrates trends in the binding behavior and correlated motions of the ARCH domain (**Table 1**). For example, the dwell time analysis of the ARCH domain’s states shows that the domain was in the “closed” conformation (“state 1”, which displayed the lowest Cy3 fluorescence) for 30% of the combined length of the trajectories. However, only 20% of the DNA binding and unbinding events occurred when the ARCH was in this state. In contrast, ~40% of the DNA binding and unbinding events occurred when the ARCH domain was in the “open” conformation (highest fluorescence state), even though this conformation was only adopted for 30% of the observed time. The results are significantly biased (p<.001 by chi-squared test) and show that binding and unbinding events are both more likely to happen when the ARCH domain is open. This conclusion aligns intuitively with the structure-functional understanding of the XPD protein (52,54,55). However, if this were a more novel system, this prevalence of binding behavior at a certain conformational state would be a vital piece of information in understanding the impact of the conformational change. **Figure 5c** shows distributions of dwell times in states 1, 2 and 3 binned logarithmically (bin size 0.4). For state 1, distributions for both, all events (n = 103) and a subset of the events when CPD-containing DNA was present (n = 31) clearly suggest at least double exponential fit. Notably, all long events (over 10 sec in duration) occurred in the presence of DNA. The ratio of amplitudes for the fast and slow exponentials for this state changed from about 90%/10% to 55%/45%. Less of a change was observed for state 2, where the ratio changed from 70%/30% to 55%/45% in the presence of damaged DNA, which no change was observed for the most open state 3 (70%/30%). This observation echoes that reported in the original study. The main difference, however, if the simplicity with which the data were extracted and sorted into distinct categories.

**Table 1.**
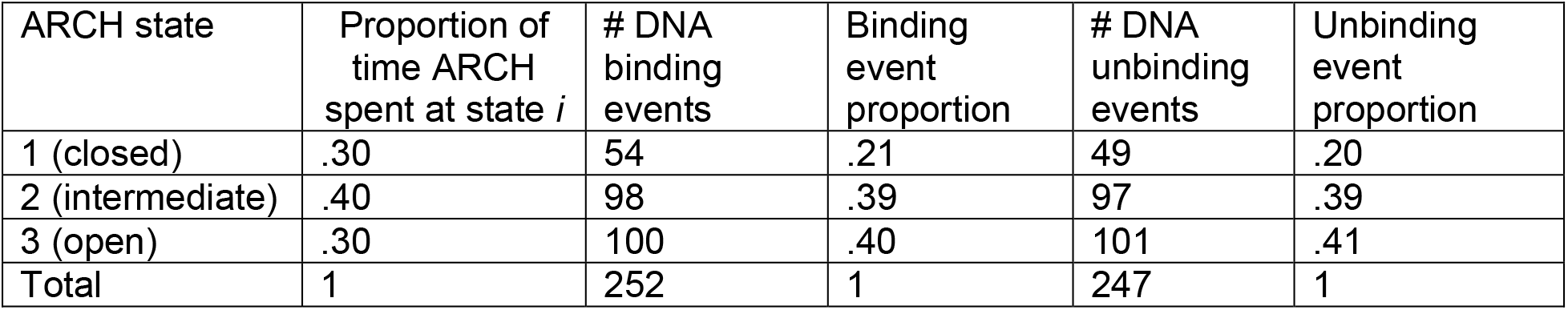
Correlation of DNA binding/unbinding with ARCH domain state

### Case Study2. Conformational dynamics of Replication Protein A (RPA)

Replication Protein A (RPA), the major single-strand DNA (ssDNA) binding protein in eukaryotes, acts as a master regulator of DNA replication, recombination, and repair processes (5). Due to the abundance of RPA within the nucleus (56) and its sub-nanomolar affinity for ssDNA (57), RPA rapidly coats ssDNA as it is exposed (see (5,58) for review). RPA has six oligonucleotide/oligosaccharides-binding (OB) folds (A-F), which are connected by flexible linkers. Four of these OB folds are high-affinity DNA-binding domains (DBD) (A-D), which can individually associate and dissociate from ssDNA within the stable RPA:ssDNA complex (**Figure 6a**). The ability of RPA to dynamically sample ssDNA binding conformation is critical for its role in directing the handoff of ssDNA to other proteins involved in DNA replication, recombination, and repair (59–61). In homologous recombination, Rad52 mediates handoff of DNA coated by RPA to Rad51, a lower affinity protein (62). In the Pokhrel, Caldwell et al. study (12), smTIRFM was used with fluorescently-labeled RPA to monitor the RPA conformational dynamics. An environmentally sensitive fluorophore, MB543, was site-specifically added to either DBD-A or D to allow for observation of their conformational states when bound to surface tethered ssDNA as varied levels of fluorescence intensity. Rad52 was also added to observe how the binding of this RPA interacting protein and mediator of RPA displacement would affect the distribution of RPA conformations.

**Figure 6:**
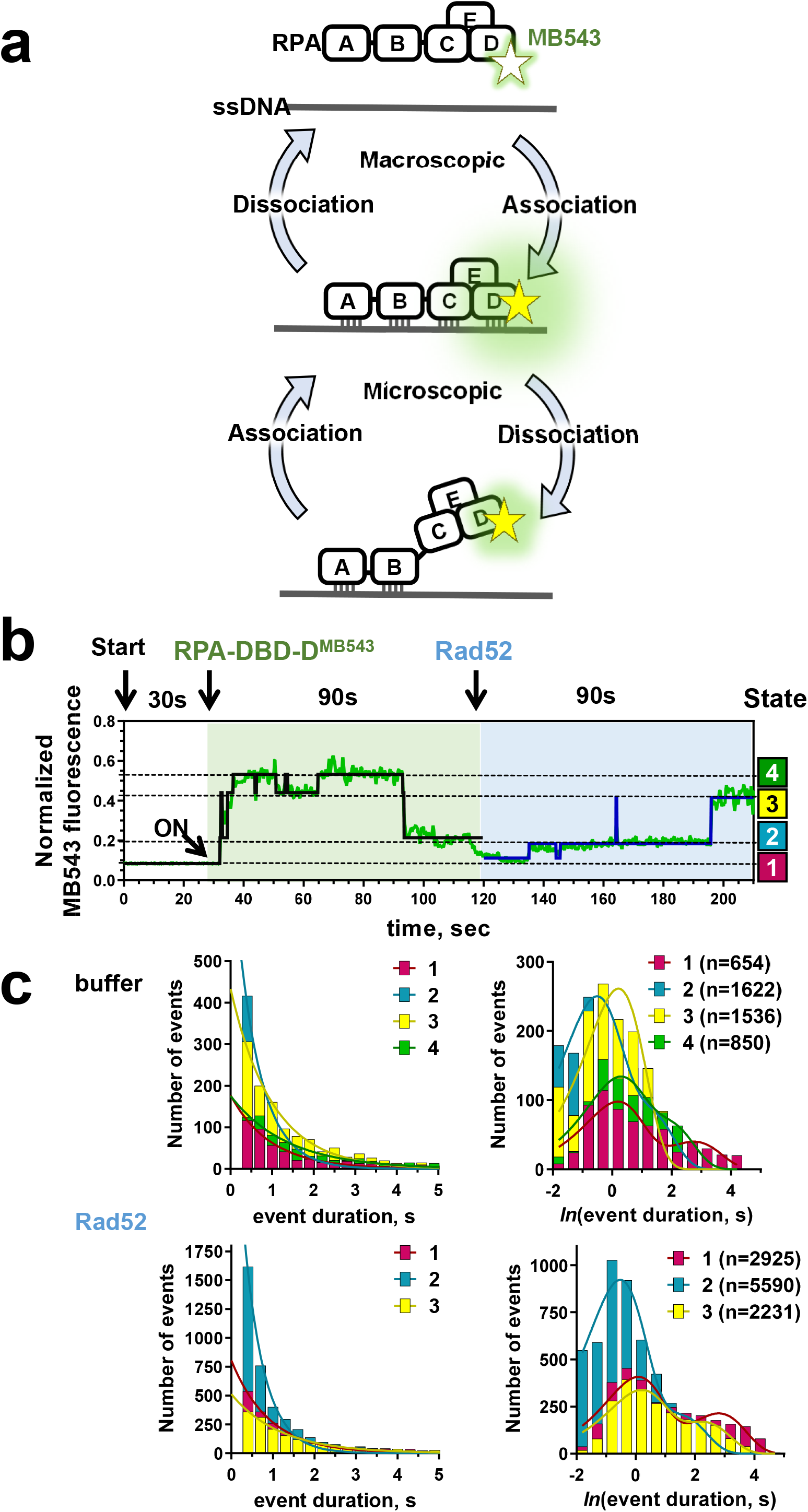
Microscopic conformational dynamics of RPA. **(a)** Conformation dynamics of RPA that results in microscopic binding and dissociation of its individual DNA binding domains can be monitored at the single-molecule level by following the fluorescence of an environmentally sensitive fluorescent dye, MB543 (green star) site-specifically incorporated into a respective DBD. **(b)** The representative fluorescence trajectory shows transitions between four states before Rad52 addition and only three states when Rad52 is present. **(c)** The dwell times for individual states are sorted with KERA and are plotted as regular or logarithmic distributions to determine the rate constants.

The selected fluorescence intensity trajectories were normalized and analyzed using ebFRET to determine a 4-state model with the four states being fluorescence intensities that correspond to different conformations of RPA (see methods). The resulting dwell times at each state were extracted using KERA. These state-specific dwell times were fit to exponential decays to yield off-rates and the percentage of events in each of the states were quantified. These values were then used to determine the effect which the presence of Rad52 had on the ability of RPA to access the four conformational states and the stability of these states. Namely, it was determined that the presence of Rad52 resulted in a loss of the most ssDNA-engaged states RPA DBD-D. This result revealed that the Rad52 mediator mechanism relies on specifically modulating the ssDNA binding dynamics of RPA DBD-D, allowing the lower affinity Rad51 to access ssDNA that was previously occluded by RPA. This is an example of the utility of KERA for analyzing a single-channel dataset to extract kinetic information.

The beta version of KERA, which was under development in the time of this study was more limited in its capabilities, lacking the custom and regex search functions. One motivation for the inclusion of those features in the current version was the desire to examine the dependence of kinetics on the past and future states of the system. With the current custom search function, the dwell times of a given state can be organized by the attributes of the preceding or following state transition. This could be useful in determining whether a system exhibits hysteresis or preferred directionality in its movements. Similar to XPD case, the RPA analysis with KERA can be readily extended to multicomponent system to separate the conformational dynamics in the presence of interacting partners labeled with spectrally distinct fluorophores. While such analysis was not necessary in the case of Rad52, which upon addition rapidly formed an RPA-ssDNA-Rad52 complex that persisted throughout the experiment, other RPA partners may form transient complexes necessitating multichannel analysis.

## DISCUSSION

KERA is envisioned as a data-processing tool for the single-molecule fluorescence experiments which operate with large data sets and explore multi-process data correlated across several colors or channels. In these, the statistical power of the kinetics and dynamics analyses are found in the large ensemble of representative events available. However, to effectively organize and study all possible events, states, and combinations of transitions, a fast and thorough approach to organizing the data of interest must be used. KERA is particularly suited to any application where discrete time trajectories are sorted by categories of transition order and transition combination.

However, fluorescence data are not the only type commonly subjected to state idealization. QuB was originally developed for studies of ion channel flows, and is now used to assign states in fluorescence studies. Similarly, the versatility of KERA, and the ability to import any time-series of integer states, means that studies outside the realm of TIRFM can also benefit from its ability to classify events. Its analysis is adaptable to arbitrarily many interacting data channels, each with any number of possible discrete states. Because of KERA’s ability to extract all dwell time data from each event, any form of analysis which makes use of dwell times may be performed on the output as appropriate. With the use of custom searching capabilities, systems may be analyzed which have patterns of arbitrary complexity, and the language of regular expressions is uniquely suited for identifying patterns with wildcard tokens, quantifiers, and logical operations. However, the user is not required to have experience with regex to use the simplified search interface, which still offers significant flexibility.

Primarily, KERA allows quick and thorough isolation of events of interest, both through an automatic general search for all event classifications, and through the optional custom search. By understanding the frequency and dwell times of the events in question, relevant properties about the biochemical system may be extracted. This increases the utility and ease of analysis for large ensembles of discretized time traces, and can reveal multi-process correlated patterns in experiments with an arbitrary number of fluorescent color channels, each modeled with any number of binding states.

## Supporting information

KERA documentation

## AVAILABILITY

KERA scripts and documentation are available in the Github repository https://github.com/MSpiesLab/KERA

## SUPPLEMENTARY DATA

Supplementary Data are available at NAR online:

Supplementary Note 1. KERA Documentation

Supplementary Note 2. Normalization Scripts.

Both documents can be also viewed in the Github repository https://github.com/MSpiesLab/KERA

## ACKNOWLEDGEMENTS

We thank members of the Spies’ and Tabei’s labs for many fruitful discussions.

## FUNDING

This study was supported by the NIH/NIGMS R35GM131704 grant (to MS), NIH/NIGMS GM081433 grant (to MTW.), University of Iowa FUTURE in Biomedicine Program (SMAT, JT), Iowa Space Grant Consortium Collaborative Grant (SMAT, MS), Iowa Science Foundation Research Grant ISF 20-17 (SMAT), University of Northern Iowa Office of Research and Sponsored Programs’ Capacity Building Grant, UNI Physics Undergraduate Summer Research Fellowship (JT), and T32 Pharmacological Sciences Training Grant NIH T32 GM067795 (to CCC).

## CONFLICT OF INTEREST

The authors declare no conflict of interest

## Notes

https://github.com/MSpiesLab/KERA

### Competing Interest Statement

The authors have declared no competing interest.

### Summary of Updates

Updated supplementary information

