## Supplementary material for "KERA: Analysis Tool for Multi-Process, Multi-State Single-Molecule Data": KERA documentation

### SUPPLEMENTARY NOTE 1
### KERA Documentation

**Background and introduction**:

KERA (Kinetic Event Resolving Algorithm) is a tool for analyzing data which consists of time-series of data at discrete states. It was originally designed for use in single-molecule fluorescence studies. In these, the fluorescence levels of individual molecules (visualized by TIRF microscopy or similar) are discretized through the use of a program such as QuB, HaMMY, ebFRET or hFRET. The resulting discrete traces have a specified number of "states". However, in many fluorescence studies, multiple reporter molecules with spectrally distinct fluorescence are used (e.g. Cy3 and Cy5). When two or more reporters are present at the same location (within a diffraction limited spot) on the slide and imaged simultaneously, they are said to be co-localized; generally, this means that the two reporters are giving information about the same chemical environment simultaneously. One detailed explanation of this type of system can be found in Boehm et al. (2016) (1), where Cy3 and Cy5 dyes were used to label the proteins PCNA and Rev1, respectively. The experiment observed the order of fluorescence appearance and disappearance in diffraction-limited locations as the proteins bound to immobilized partner proteins. Distinguishing the protein species was possible because the fluorescent dyes had different colors. These distinct signals are referred to as *channels* within KERA; in this case, the two channels corresponded to two different binding partners. In other cases, such as the study of XPD mentioned in the main text, a channel might be dedicated to fluorescent reporting of another aspect of the system, such as the conformational state of a protein domain. KERA supports any number of channels, and any number of states within each channel. KERA's purpose is to take in discretized trajectories and organize them in a method which makes it possible to see the transitions, interactions between channels, and kinetics of the events contained within. Whether the user searches for the default set of patterns, defines their own using the convenient user interface, or leverages the limitless flexibility of the raw regular expressions input, KERA is built to find multichannel interactions of interest in discrete-state time traces from any source.

**Pictorial overview:**

**Figure S1** shows three examples of fluorescence trajectories, along with the molecular activity that might produce them. In a real experiment, the brightness of the single molecules would be a noisy signal, and then some idealization process would be needed to assign an integer-level state to each time point in the trace (as seen in **Figure S2**).

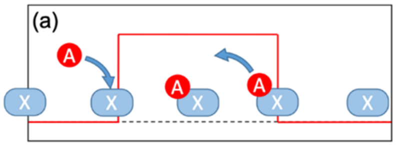

a) a fluorophore binds to an immobilized molecule, resulting in a spike of fluorescence which lasts for a certain amount of time. The idealization would look something like [1 1 1 1 … 2 2 2 …. 1 1 1 ].

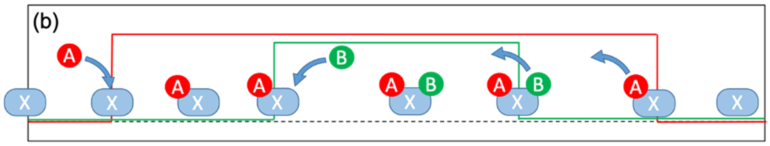

b) a two-channel scenario, where red and green fluorescence are both measured from the same location

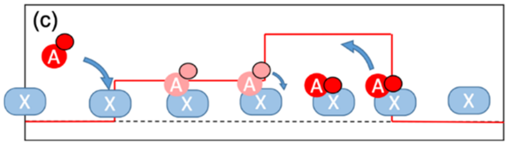

c) a fluorophore binds, resulting in some fluorescence, and then a conformational change results in a brighter fluorescence. [1 1 1 … 2 2 2… 3 3 3 … 1 1 1 ]. A 3-state model.

**Figure S1**

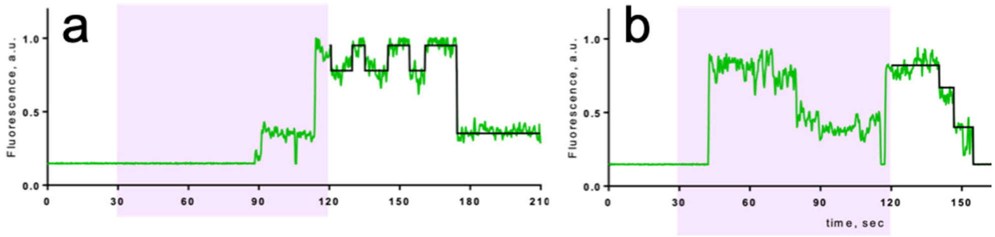

**Figure S2**: Examples of idealizing a noisy trace to a 4-state model

**Figure 2**

The discrete traces are long lists of integers in files. In multi-channel setups, there will typically be a file for each channel, and some method of making sure that the files that contain the colocalized pairs match up with each other. The first thing KERA does is to locate all of the transitions (changes of state) within each trace (or colocalized set of traces). This allows it to condense the traces from a string of integers, one per time point, into an ordered list of the configurations which the system achieves, paired (in a different variable) with the time points associated with the transitions. So [1 1 1 1 …. 2 2 2 2 …. 1 1 1 1 ….] becomes [ 1 2 1 ]. For multi-channel systems, the channels get their own columns within the array, and will be displayed for simplicity as such: [1 1 ; 2 1 ; 2 2 ; 2 1 ; 1 1 ] matches panel b of the first figure above (with the convention that the red line is channel 1 (the first number in each pair)).

The word “configuration” refers to the combination of states at a given moment for a given trace set. For single-channel data, the current configuration *is* the current state. The distinction becomes important in multi-channel data; if channel 1 is at state 3 and channel 2 is at state 1, then the configuration of the system is [3 1].

The word “event” refers to a series of configurations; in KERA, the user can search for events and the set of instances of that series of configurations within the data will be compiled for easy reference. This is called defining a classification. KERA has tools which allow the creation of classifications which have wildcards or variable length. These are described later.

**Getting KERA started:**

Download the code from Github: https://github.com/2718xyzu/Spies-lab
If you don’t have git, Github, or any other source control on your machine, “Download as Zip” is fine; extract the zip’s contents to somewhere that your MATLAB can access. When you open MATLAB, you should change the search path to include the sourceCode folder (you can do this by using the “browse for folder” button in the path toolbar). Alternatively, you can open the main code file: openKERA.m, and run it; you should be prompted to “change folder”; this will change the search path automatically. Further details on source control can be found in the appendix to this guide.

Running the openKERA script presents the user with a blank interface; the first step is to import data. Data is currently supported for import from ebFRET, HaMMY, hFRET, and QuB programs. The conventions for each type of import are listed below; it is important to note that, for multi-channel data sets, the naming of the individual files is sometimes critical to making sure the proper traces from each channel are eventually paired correctly. Advice on file naming conventions is given later.

Before importing, the user is prompted to indicate the number of channels in the data set and the number of states in each channel. Once set, the number of channels cannot be changed, though the number of states in each channel can be edited by viewing the data and editing the states there. The number of channels is the number of colocalized, distinct reporters, which interact with each other. The number of states in each channel are inputted in a comma-separated list. For example, if channel 1 has a 3-state model and channel 2 has a 2-state model, the input for the second box would be 3,2. Some examples of various types of input files are included in the KERA repository.

The user is also prompted to select a time step for the program; in general, this is equal to the duration of each frame from the data, but in the case of QuB it may be something else (like .001, for the millisecond resolution which it seems to output its time lengths in). 0.1 is a common value for microscope movie resolution.

- ebFRET (3,4): after analyzing the traces from one channel in ebFRET (available from https://github.com/ebfret/ebfret-gui), export the desired state model using the menu and selecting “Single Molecule Dataset (SMD)”. In this way, an SMD should be created for each channel. The order in which the traces were imported to ebFRET impacts how they are ordered in the SMD, and thus how they are paired in later analysis. You can check to see that files were paired properly by debugging the program in the script “packagePairsebFRET” (located in the “functions” folder) near the end and looking at the “filenames” variable.
- QuB (5): QuB’s file naming conventions are particularly complex, and this method of import has the least support in KERA, generally. All file names must be of the form # tr#*[suffix].dwt, where the first # symbolizes a single digit, “tr” refers to the literal string “tr”, #* refers to any string of digits which identifies the trace, and [suffix] refers to a string

(which begins with a space, special character, or letter) which identifies the channel which the trace came from. Valid examples of a file name are

  - 1 tr123 Cy3.dwt and 1 tr123 Cy5.dwt are a valid pair
  - 2 tr123 C3.dwt and 2 tr123 CY5.dwt are also a valid pair, distinct from the ones above because of the initial "2", but can still be sorted into the correct channels (the channel suffix algorithm is purposefully coded to allow typos and deviations, since the data set originally used to test the QuB import function had such typos).
  - tr123_chan1.dwt and tr123_chn2.dwt are valid but may not be distinguishable from the first example given because they do not have an initial number (which defaults to assigning them a "1").

  QuB data must be imported by first placing all dwt files into the same folder. To my knowledge, that is how QuB outputs its files; if it is not, you can just make a copy of your existing dwt's and place them into one folder. QuB is not a preferred datatype because, though it was what KERA was first applied to, no one I have talked to since has used QuB or given advice on how QuB is typically used. If you would like KERA to be better-prepared to handle QuB input, please let me know and I'd be happy to have a conversation.

- HaMMY (6): HaMMY is freely available for PC's from http://ha.med.jhmi.edu/resources/#1464200861600-0fad9996-bfd4[3]
  It helpfully places all output files into the same folder as the import files, so when prompted to select the folder for each channel, it will just be the folder which contains all the output from that channel's analysis in HaMMY.
- hFRET (7): this is also available as a github repository at https://github.com/GonzalezBiophysicsLab/hFRET. After running the hFRET analysis on your traces, you must click the "save" button. This feature is still somewhat in development, and only works on analyses where there is one "dimension" of the HHMM and 1 "static subpopulation", leaving the model with a simple integer number of levels and no hierarchy of levels.
- Raw data: this allows the user to import data which has been saved as a MATLAB variable, specifically a cell array in a certain format. If your data is already discretized, and you aren't using any of the listed discretization software, rearranging it into this cell array format and saving it is the easiest way to get it into KERA. See the appendix for formatting requirements.
- If there is another form of discretized data which you think should be importable, some comments in the scripts "openKERA.m" and "Kera.m" will help you set that up.

Once your data is imported, you can go to the analysis tab and "view data" (which gives you the ability to see each trace's discretization and, if available, the raw data behind it, for all channels at once). If you're less interested in viewing the data (possibly because you already saw it in the ebFRET window), you can skip the next few pages and proceed to the "Run Analysis" section.

Within this interface, buttons appear at the bottom of the figure to allow navigation. The filename, if available, is displayed at the top. A legend reminds the user which color corresponds with each channel (the legend can be moved by dragging it to reveal data which it might be obscuring). Numbers at the right of the screen identify the numerical value of each discrete state, at the level of the corresponding state. This is necessary because some discretization programs are not careful when it comes to assigning the states numbers which correspond strictly with the fluorescence levels; in some sets, "state 3" will have a lower average value than "state 1" which is itself lower than “state 2”. When this happens, you may consider using some of the tools included to swap the order of the states.

However, an important caveat: discretization software is not perfect, but it is based on thoroughly-researched methods and robust statistical analysis. The tools in this viewer are to be used sparingly and judiciously, since the human eye is good at seeing patterns which aren’t there. However, if changes are justified, it is the hope that this simple interface makes some of them easier. The tools are as follows (all of these are defined by the callback code of plotDisplayKera.m and the interface itself is built using the function KeraSelectUi):

- “Discard Trace”: click this button to remove the current trace from the analysis; you can restore it by viewing the trace again and clicking the “reset” button. “Reset All” restores all discarded traces at once, in addition to reverting them to their original form.
- “Next trace”: move forward through the traces
- The drop-down menu will move to the indicated trace index. Any changes made to the current trace are automatically saved when navigating to another trace.
- “Threshold”: this command opens up a new window which displays the thresholding interface. The histograms display the fluorescence values which are associated with each state. This process is best explained with an example. In the figure below, the user has decided that some of the dwells in state 2 (orange histogram) are a continuation of the baseline (blue histogram) which due to instrument noise happened to have a higher fluorescence value and were subsequently assigned to state 2. To rectify this, the user has selected the following options (see figure below):
    - Channel: Select which channel to be edited (the channels are independent of each other and must be edited separately; channel 1 is the one with the three-state model which we are interested in modifying in this example)
    - Below/Above/Between: this dropdown specifies which data points will be re-written. The numerical boxes allow the input of a number; in this case, the user wants to edit all dwells which have an average fluorescence *below* the value of 0.3 (corresponding to that small peak in the orange histogram).
    - Absolute/Relative value: changing this box changes how the values of fluorescence in each trace are treated. “Absolute” takes into account only the actual value of fluorescence; “relative” normalizes each trace (placing the 1$^{st}$ percentile value at 0 and the 99$^{th}$ percentile value at 1, to lessen the effects of noise spikes) before analysis. This would allow the user to specify that all dwells with a center above the “midway” point of each trace should be assigned to state 3, for example. In the current example, the user is solely interested in the absolute value of 0.3.
    - Currently at: the options are “any” and the numbers corresponding to the states in the model. If “any” is selected, the above thresholding criteria will be applied to all dwells, regardless of their current state value. Here, the user is interested in changing only the values which are currently state 2 (though, because the only other dwells matching the above criteria are already state 1, it does not matter in this case)
    - To State: The state to which the thresholded dwell states will be changed to; an integer greater than 0. This can even be a state which does not currently exist in the model. This is useful when it is desired to swap the identities of two states. If states 3 and 2 need to be swapped, threshold all state 3 dwells to state “4”, then threshold all state 2 dwells to state 3, finally change all state “4” to state 2, and

the swap is complete. The user in this example has set the state of interest to be 1.

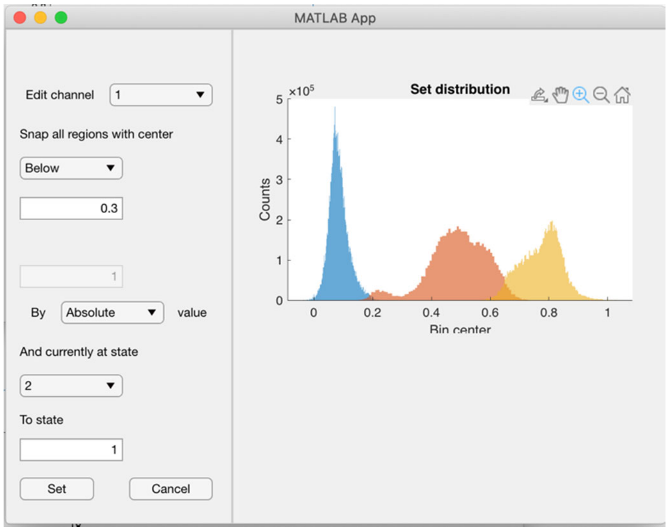

After the “set” button is pushed, the thresholding function may be used again to check the histograms; below is the result:

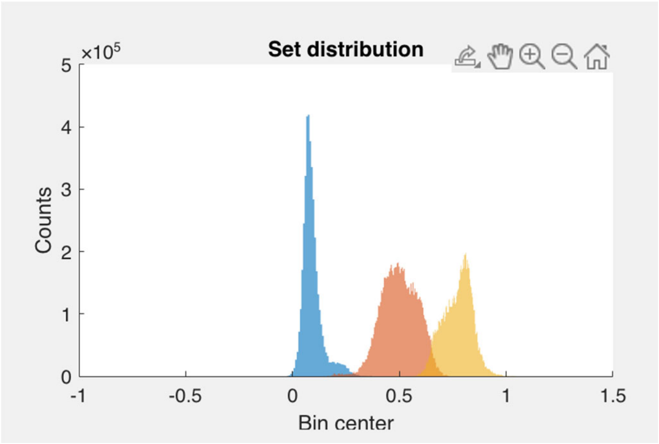

- Once a thresholding has been applied to the data set, a new button appears which says “Undo Threshold”. This button will only undo the most recent thresholding; if a second one is applied, the condition of the data set before the penultimate thresholding is no longer retrievable (though the “reset all” button still exists for the reversion of all traces back to their very first, original state).
- Manual Assign: a useful but potentially dangerous tool (from a data integrity standpoint). Clicking this button allows the user to use the brushing tool (click and drag) to highlight data points. Using shift+click will allow multiple regions to be selected simultaneously. When the “Assign brushed” button is selected, the user is prompted by dialogue boxes to specify a channel to edit and a state to manually assign the brushed data to. The result is that all discrete data points of the specified channel will be re-assigned to the

specified state. (Raw data points, if selected, are simply ignored). At any point during the brushing, the "cancel" button can be used to stop the manual assignment.

- Reset: always reverts the current trace to the way the data was upon import. All changes made to that trace individually will be reverted.
- Reset All: the entire data set is reverted to its original form. The previous two buttons can be "undone" by closing out the view window without saving; this causes all changes made within the last view session (including the "reset" action itself) to be deleted. This is why it is a good idea to go through multiple view sessions, starting with more minor changes, saving those changes, and only then open up view sessions to experiment with more drastic changes; this way, closing the window without saving will only undo the most recent changes.
- Save and Exit: this button saves all changes that have been made in the current view session. If you have already run an analysis and are viewing results, you must run the analysis again in order for the changes you made to take effect in the results you see.

For all of the above tools, Clicking "Run Analysis" will never clear out the custom event searches you have already done (though it may add new event classes depending on what it finds in the edited data). It will use the previously-added search classes to go through the (now edited) data.

**Notes and troubleshooting you might encounter while viewing data**:

- Closing the viewer window exits the viewer and saves all of the changes you have made to the data using the aforementioned tools; it also allows you to go back to the main interface and run analyses (always close the viewer before interacting with the other interface).
- The trace viewing interface relies on the "uiwait" command to pause progression of the program while the interface is open; this means that some other matlab processes (like editing code, executing other programs from the command line, etc.) may not work properly while the interface is open.
- Underscores in the filenames will be interpreted to create subscript characters when Matlab parses them.
- If the traces on the screen don't appear to be the same length, they may have been paired incorrectly (the traces you see are not colocalized in the original data), and you should check the import function.
- The threshold function only moves whole, unbroken state dwells; it cannot split an existing dwell into multiple pieces. This is because it measures the average value at which each contiguous state region exists in each trace before deciding whether to re-assign that region's state. It does not look within an existing dwell region to find places where it might re-assign only a portion of the points to a new state. That is not KERA's job, because it requires an understanding of Bayesian statistics and/or Markov models which is already present in things like ebFRET and HaMMY.

**Run/Refresh Analysis:**

This is the "big button" of the Kera code, as it were. The command which runs the analysis for which KERA was envisioned: a search of the data for all "interesting" patterns contained therein. Of course, "interesting" is a subjective term, and while the default analysis tries to find patterns of a specific form, running the "Custom Search" after the default analysis is often necessary for the researcher to really find what they're looking for. Of course, if you're very sure what you're looking for, you can simply run the "Custom Search" on that pattern (or run a few custom searches for multiple patterns) and don't bother with the initial run of the default search; more information on that can be found in the "custom search" section.

Whenever the "Run/Refresh Analysis" button is pushed, any previous searches which had been run are copied and run again on the new data. So, if new data had been imported, or the data had been modified using the "view data" interface, the updated analysis will use the same search parameters on the new data by default. This is why the button is labeled "Run/Refresh". However, in addition to the old searches (if any) being used again, the user is presented with questions about what kinds of search they would like to run.

The first question asks if the user would like to search for all event classifications which start and end at the "baseline" configuration. Typically, the baseline is set to the "default" configuration of the system (such as the configuration in which nothing is bound, or all species are in their most stable conformation), so the regions *in between* baseline states are where the interesting events (bindings, conformational changes) are often events of interest. More information on setting the baseline can be found under the "settings" section, but a baseline of state-1 signals in all channels is the default which is set when KERA is opened.

The second question asks whether the user is interested in a compilation of all the possible system configurations. This is useful, kinetically, for learning how long different configurations tend to last. So, if the system has two channels (each with two states), this analysis would produce a search of all occurrences in the data of [1 1], [1 2], [2 1], and [2 2] configurations. The resulting dwell times (along with the other data about the search terms which is described more in detail below) would be displayed among the results.

If the user selects to find all possible configurations, the user also has the option to find all possible single-channel transitions within the dataset. Examples of single-channel transitions are: [1 1; 1 2] and [1 1; 2 1]. In both cases, only one channel changed in the course of the transition. This can be useful because, in some systems, the dwell time of the initial [1 1] state might be affected by the state to which it transitions.

Of course, the above options are meant to give a wide picture of the data imported; if the user has a specific set of transitions or event they want to study, it is generally easier to run a custom search. In a large data set, or a system with many possible configurations, the above options can produce dozens or hundreds of distinct classifications. For ease of analysis, they are organized by prevalence (the number of times that pattern occurs in the data) with the most common ones first. And even large data sets can be seen somewhat at a glance by going inside the kera variable in the workspace and finding the "output" field—sorting the event classifications by the meanLength or searchMatrix fields might help organize the results more conceptually as well.

Speaking of the "output" field, it is worth describing here what data is contained in this variable after the analysis is complete. kera.output is the main location of all results from the analyses which have been run on the data. When the analysis is run, a copy of the variable is made and placed into the base workspace (called 'analyzedData' to distinguish it from the original), since not everyone is comfortable navigating the workspace inside of an object class like Kera. Changes made to the kera.output variable do impact the data displayed on the frontend; changing how kera.output is sorted changes the order in which results are displayed on the main interface (editing analyzedData does nothing; the changes don't back-propagate). kera.output is a structure, with the following fields, each of which is recorded for every event classification (event search) on its own row:

- searchMatrix: the numerical representation of the states and transitions which are searched for. Conventions are explained in more detail later. In brief, the matrix has a column for each channel and a row for each dwell; NaN stands for a wildcard (any state) and a row of Inf denotes a series of one or more dwells. So, [2 1; Inf Inf; 2 1] finds all places in the data where the system starts at channel 1-state2 and channel 2-state1, goes through some number of transitions, and returns there again.
- expr: searchMatrix turned into a string using the mat2str function. This is to allow the function 'unique()' to run on the elements.
- count: the number of times events matching the search string occurred in the data
- meanLength: the mean length of the events, in units dependent on what was entered for the time-interval. If the frame rate is 10 frames/sec and the time-interval was entered (correctly) as 0.1, this column will be in units of seconds. Because dwells at the beginning and end of traces have indeterminate length, the dwell time of the first and last configurations of a given search are *not* counted; the search [1 1 ; Inf Inf; 1 1 ] only counts the time spent *in between* the baseline configurations. That is why searches for an individual configuration must be worded as [NaN NaN; 1 1; NaN NaN]; to get an accurate picture of how long the system spends at the [1 1] configuration, the search has to count only those dwells where some other dwell happened before and after the [1 1] configuration. The dwell times for a search such as [1 1 ; 1 2 ; 1 1] only represent the length of time the system was at [1 2].
- randomnessParameter: as described in Schnitzer and Block (1995) {schnitzer block 1995} for a process which has independent, first-order rate-limiting steps, the randomness parameter *r* is a dimensionless quantity which can be used to estimate a lower bound on the number of rate-limiting steps. If $\tau$ is the set of residency times in a state or series of states, $r = (\langle\tau 2\rangle-\langle\tau\rangle 2)/\langle\tau\rangle 2$. A single-step first-order reaction would result in a randomness parameter near unity; reactions with p rate-limiting steps produce lower values of r, and a lower bound on the value of p may be approximated by 1/r. The variability of r depends on the number of events in the set under study, and iterated random sampling of the list of event lengths (available in the TimeLengths field) should be used to understand this variability before conclusions are drawn based on the reported value of *r*.
- EventList: an explicit listing of the events which make up the classification. For classifications without any wildcards, all entries in this list are identical to searchMatrix.
- TimeLengths: the dwell time of all instances of the event in a list (see meanLength above for conventions of measuring dwell time).
- table: a comprehensive set of details about every event in the classification. It contains the following columns and every instance of the classification gets its own row:
    - Events: same as EventList
    - Total_Duration: same as TimeLengths
    - Time_Points: the points in the trace which mark the beginning and end of every dwell in it; this list *does* include the first/last states in the search term
    - Delta_t: the time difference between the points in Time_Points
    - Time_First: the first element of Delta_t
    - Time_Last: the last element of Delta_t
    - File: the file (if provided) in which this instance of the event occurred
- excel: a field made specifically to enable copy-pasting of the data into another spreadsheet program. This variable is only created if the event classification has an

explicitly constant number of dwells (no Inf wildcards and all instances are the same length). With this criterion met, the created variable is a rectangular array with a row for each instance of the event classification; the number of columns is equal to the number of dwells in the full search parameter. The elements of the array are the dwell times of each step in the event for that particular instance.

**Custom Search:**

At any point after importing data, the user may choose to add a “row” to the output structure by specifying a search term, a custom classification that KERA will search for within the data. When the “custom search” option is selected, a new dialogue box will appear which allows the user to easily specify the pattern they would like to look for. The user is presented with dropdown menus which list the states available in each channel; making a selection updates the first configuration of the search term; to add sequential configurations, click the “add to search” button; to remove the last configuration, click the “Remove Previous” button. The N-state wildcard refers to a series of configurations which has any length (or no length at all). For example, in a one-channel data set, it might be used in the following way: the user is interested in all the ways in which the system evolves from state 1 to state 4. They would use the custom search function to search for “[1 Inf 4]”. This would create a new row in the kera.output variable which would include, for example, the event “[1 2 4]” and “[1 3 4]” and “[1 2 3 2 3 2 3 4]”, as well as simply “[1 4]”. By going to the output structure and selecting the entry under “table” for the appropriate row, the user would be able to see each of the instances of the above patterns, as well as sort them by duration or by classification. If one of the classifications appears promising for kinetic analysis, another custom search could be done for any specific event type (such as [1 2 3 4]).

It is important to keep in mind that, for any search, the “total time length” which is used in the kinetics fittings and displayed in the meanLength field does not take into account the dwell time of the first or last configurations in the search. This is for two reasons: first, many searches are focused on finding the dwell time of an interval between two configurations (for example, for the “off time” of some binding, which would be found by searching [2 1 2], it is the dwell time of the “1” state which the user is interested in). Second, the dwell time of any configuration at the beginning or end of a trace is not known (being cut off by the edge of the data), any measurement of the dwell time of a given configuration must ensure that at least one configuration (of any type) happens both before and after it. So, a search interested in the dwell time of individual state-1 dwells would be phrased as “[NaN 1 NaN]”, where the “NaN” wildcard matches any state. This will match only (and all) the state-1 dwells which are not at the beginning or end of a trace.

**Table S1** below shows some possible custom search sequences; some of the search sequences are for one channel data, and some for two channel data.

**Table S1**

| Search Sequence: | Explanation | Possible matches; timeLength measures the bolded portion; underscores are wildcards |
|---|---|---|
| [1 2 3] | An explicit set of transitions; the meanLength field records only the length of time spent at “2”. | [1 **2** 3] |

| | | |
|---|---|---|
| [NaN 1 2 3 NaN] | The same set of transitions, but with the restriction that they not occur at the beginning or end of a trace; also, meanLength and kinetic data are accessible for all three of the dwells. | [_ **1 2 3** _]<br>For example:<br>[3 **1 2 3** 2]<br>[3 **1 2 3** 1]<br>etc. |
| [1] | This is not a recommended search string | [**1**] |
| [NaN 1 NaN] | This will match all instances of state-1 which can be kinetically analyzed (are not at the border of the trace); meanLength measures the dwell time of state 1 alone. | [_ **1** _]<br>For example:<br>[2 **1** 2]<br>[2 **1** 3] |
| [1 Inf 4] | All evolutions of the system starting at state 1 and going to state 4; meanLength measures the time spent *between* states 1 and 4. | [1 4] (timeLength of 0)<br>[1 **2** 4]<br>[1 **2 3 2 3 2 3** 4]<br>etc. |
| [NaN 1 Inf 4 NaN] | Same as above, but the time lengths recorded include the time spent *at* states 1 and 4 as well as the time between them. | [ _ **1 4** _ ]<br>[ _ **1 2 4** _ ]<br>[_ **1 2 3 2 3 2 3 4** _] |
| [NaN 1 NaN Inf 4 NaN] | Same as above, but there must be at least one intervening state between states 1 and 4. | [_ **1 2 4** _ ]<br>[_ **1 2 3 2 3 2 3 4** _]<br>etc. |
| [NaN 1 NaN NaN Inf 4 NaN] | Same as above, but now there must be at least *two* intervening states | [_ **1 2 3 2 3 2 3 4** _]<br>etc. |
| [NaN NaN;<br>1 2 ;<br>NaN NaN] | (Switching to two-channel examples now, search strings are listed on multiple lines for clarity); an explicit search for the [1 2] configuration. | [_ _; **1 2**; _ _] |
| [NaN NaN;<br>1 2 ;<br>2 NaN] | A subset of the above, but only including the events which are followed by a transition in channel 1. | [_ _; **1 2**; 2 _] |
| [ 1 1 ;<br>Inf Inf ;<br>1 1 ] | One of the “baseline” searches; all events which occur between instances of the [1 1] configuration. | [1 1 ; **2 1** ; 1 1]<br>[1 1 ; **1 2** ; **2 2** ; 1 1]<br>etc. |
| [ 1 1 ;<br>Inf Inf ] | This is not a valid search item; n-state wildcards must be followed by another search configuration so that the search engine can terminate the event. | (no matches) |
| [NaN NaN;<br>1 NaN;<br>NaN NaN] | This is not a recommended search, mainly because the user cannot know whether it was Channel 1 or Channel 2 which transitioned next; if the user is interested in looking at the behavior of channel 1 independent of anything happening in channel 2, the user should just open a KERA session which is single-channel and only import the channel 1 data | [_ _ ; **1** _; _ _ ]<br>So, for example,<br>[_ _ ; **1 1**; 2 1 ]<br>[_ _ ; **1 1**; 1 2 ]<br>would both be included, but in the latter case, the state-1 dwell could extend beyond the time recorded (**1 1**) |
| [ 1 NaN;<br>2 NaN;<br>Inf Inf; | A possible workaround for the above problem. This measures the “on time” of | |

| | |
|---|---|
| 1 NaN] | channel 1 (the time which it spends away from state 1 after transitioning to state 2) |

The KERA interface post-analysis:

Although looking through kera.output (or the analyzedData variable) are good ways to see a lot of data at a glance, the KERA interface is designed to help visualize certain aspects of it. For example, each row in the output variable is displayed visually by the top panel; the user can scroll through the rows using the arrow buttons on the interface. Wildcard states are denoted by a blank space, and n-state wildcards are denoted by a blank space with an ellipsis superimposed on it. Below is a screen capture of the interface:

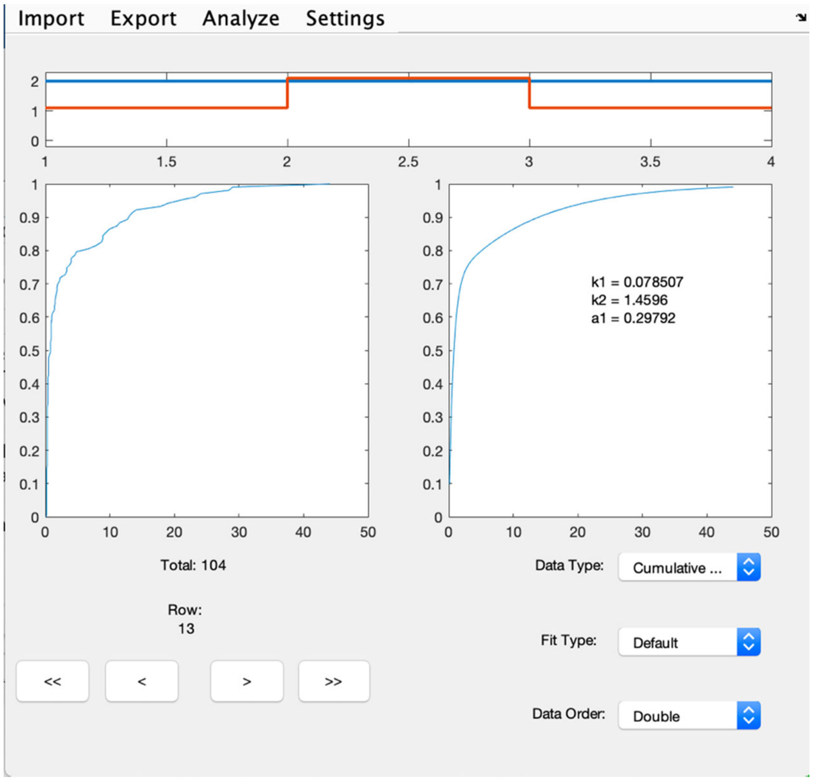

The top panel shows that the current classification of events being viewed is “[2 1 ; 2 2 ; 2 1]”. The blue trace corresponds to channel 1 and the orange to channel 2; this pattern, of following the MATLAB standard color order, is used in place of including a legend on the small display.

On the bottom of the screen are the navigation controls; currently displayed is row 13 (the data contained in the 13th row of the output variable), and the “total” display shows that 104 instances of the displayed configuration exist in the data. The arrow buttons can be used to move to the next/previous rows, as well as to the first or last rows.

The other displays are a graphical representation of the dwell times of the event in question. As mentioned above, the dwell time recorded for this event classification does not include the first or last states; here, the dwell times refer only to the time spent at the middle [ 2 2 ] configuration. Those dwell times (104 of them) are collected into a visual representation in the left panel and an idealized curve-of-best-fit in the right panel, the nature of which is determined by the dropdown boxes on the lower right.

- Data Type: Histogram or Cumulative Distribution. Fitting an exponential to the bin counts of a histogram is common when conducting a kinetic analysis, but for events which are relatively rare (~100 events or less) the cumulative distribution may be preferable. It is not dependent on bin size, though adjacent data points are no longer independent. In Figure 3, the cumulative distribution is shown (indicating, for example, that 80% of the dwell times are shorter than 10 s).
- Fit Type: "Default" uses the typical exponential fit for the data, while "Logarithmic" uses the strategy of taking the natural logarithm of all time values before fitting (using a different fit model, of course). Details about logarithmic fitting can be found in Boehm et al.[2] and Szoszkiewicz et al.[4]
- Order: "single" fits the data by assuming that the times are distributed according to a single-exponential distribution. "Double" fits the data by assuming that the times are distributed according to a double-exponential distribution (with a PDF of the form $f(t) = a*exp(k_1t)+b*exp(k_2t)$ ).

The function responsible for fitting a given group of time data is getFitHistogram.m.
It is important to note that KERA's methods of fitting dwell times are rudimentary; a more complete analysis would involve a bootstrapping process to get confidence bounds on the parameters and a more careful binning strategy for the histogram fits. However, the fits given should provide some qualitative insight into the approximate size of the rate constants involved.

All of these fitting strategies rely on the assumption that the dwell times are exponentially distributed. Of course, searches which result in dwell times which are one configuration long (such as "[NaN NaN; 1 1 ; NaN NaN]") will often be amenable to this analysis, but many searches are specifically for event types which contain multiple configurations sequentially. The total dwell time for this type of event will not be distributed exponentially (the sum of independent random variables from exponential distributions is a gamma distribution). If the kinetic data of any particular step within these events is desired, the dwell times for specific steps in the set of events may be displayed. This is done by going into the settings tab and selecting the "Toggle intraevent kinetics". Now, whenever a classification is viewed which has multiple configurations as part of the event (such as "[NaN NaN; 1 1 ; 1 2 ; NaN NaN]"), the user is given the option to select which part of the event they would like to view kinetic information for. A yellow highlight within the top panel will show which part of the event's dwells are being analyzed.

Other outputs: the stateDwellSummary
This is a variable which is outputted into the workspace and also exists as a field of the KERA object. This was included because there are times when the researcher is interested in knowing some information about the traces' channels independently of each other. For this example, we'll say that the baseline is set to [1 1] (state 1 in both channel 1 and channel 2), though the base state could be anything. Each row corresponds to a channel.

- dwellTimes: a cell array with as many cells as there are states in the channel; each cell contains a list of the lengths all contiguous regions of that state in that channel. However, it excludes the instances of states which were cut off by the beginning or end of a trace. So in a trace which starts with 10 seconds of state 1, then 10 seconds of state 2, then ends with 10 seconds of state 3, only the second cell array will have a value of 10 appended to it.

- meanDwells: the mean value of the dwell times contained in each of the cells in the dwellTimes array
- dwellTimesWithEdges: it contains all of the same data as dwellTimes but also contains the states which were cut off by the end or beginning of a trace. If the researcher is interested in calculating the proportion of time which the system spent at a given state, regardless of where they occurred in the trace, this is the data to use. In the example given in the above bullet point, all three cell arrays would contain an entry for 10 s from that trace.
- meanDwellsWithEdges: again, this is just the mean of the dwellTimesWithEdges arrays, but it allows the researcher to quickly calculate the total time spent at each state. Just multiply each mean value by the length of the corresponding cell array held in dwellTimesWithEdges. The total of these should also equal the combined runtime of all non-excluded traces in the dataset.
- timeBeforeFirst: This contains all of the dwell times where the channel in question was at the *baseline* at the beginning of the trace. In other words, it contains the time spent before the first "event" (or transition from baseline) occurred.
- timeAfterLast: a similar field, though this one records the amount of time spent at the baseline state for the given channel at the *end* of the trace. In either of these last two fields, there may be as many entries as there are traces (if the baseline state is very common) or there may be very few.

The below image is an example of what the stateDwellSummary looks like for a two-channel example, where the first channel has three states and the second has two states:

| dwellTimes | meanDwells | dwellTimesWithEdges | meanDwellsWithEdges | timeBeforeFirst | timeAfterLast |
|---|---|---|---|---|---|
| *1x3 cell* | [7.4846,6.6608,5.7077] | *1x3 cell* | [23.9730,12.3185,10.27... | *1x17 double* | 5.3000 |
| *1x2 cell* | [22.5441,7.5928] | *1x2 cell* | [25.6716,8.8529] | *1x48 double* | *1x43 double* |

In channel 1, seventeen traces begin with the base state, and only one ends with a base state dwell (which lasted 5.3 seconds).

Regex (Regular Expressions) analysis:

See appendix for technical details about this advanced search method, which allows for patterns with greater flexibility to be defined.

Open in cftool:

This option under the "Analysis" menu allows the data displayed in the left panel to be opened in Matlab's built-in interactive fitting environment, cftool. The cumulative distribution values (or histogram bin heights and positions, if the histogram option is chosen) are exported to cftool in a new window, and also saved to the base workspace as variables. The user can then fit the distribution to exponential models or custom equations and get statistical information about the fits.

**Exporting:**

The easiest way to export a given set of data is to find it in the structure, highlight it, and copy-paste it into a spreadsheet application. Then kinetic data may be extracted using whatever established method the researcher is familiar with. A more automated export can be accomplished by selecting the "Export→ Analyzed Data→csv" option, which will create separate csv files containing the information from the "table" field of each row and save them all in the same folder. The stateDwellSummary can also be exported in this way.

**Saved Sessions**:

In the course of analysis, the researcher has the ability to save the entire KERA interface, and all changes made to the data, in a convenient package. This is done by selecting the "Export→Save Session" option. This creates a MATLAB variable which stores all aspects of the current analysis in such a way that any version of MATLAB with KERA installed (and on the path) can open it. It is important to note that the source code behind the KERA interface is not saved with the variable; it still needs the code files from MATLAB in order to run. For this reason, if you try to open a saved session which was made by an outdated version of KERA, or someone who has edited the source sends you their version of a saved session, there is no guarantee that it will behave in the same way. This is why, if (or when) updates to the KERA code are made, special consideration will be made to keep old saved sessions still operable, and under-the-hood changes will be explained clearly in the Github log.

When a saved session is opened, a new KERA window is opened, and a new kera variable will appear in the workspace (with a number which corresponds to the figure window which it occupies, to avoid overwriting any existing Kera variables, e.g. kera2 or kera4). Both the new and old KERA's (and any subsequent saved sessions which are opened) can be manipulated independently, though it is worth keeping in mind that some editing functions (like the view data function and thresholding) affect the system-wide MATLAB timing and wait functions, and so unexpected behavior may result if two separate KERA's each have these open.

A saved session can also be created by typing into the command line "[kera variable name].exportSPKG", which is useful if the window has been closed but the variable is still available in the workspace, or if the session saved itself as a workspace variable during a MATLAB crash.

**Settings:**

Setting the baseline state: the default behavior of KERA is to search for the types of "events" that happen; in many binding studies, this means episodes of binding, which are separated by "off times" of no activity. To demarcate "events", the code assumes some kind of baseline, which corresponds to a configuration which the system departs from and returns to in between the events. So, for a simple binding study where high fluorescence in either channel means that a binding has occurred, the baseline is just a state-1 signal in all channels. Say there are two channels; from the [1 1] configuration, the system might evolve such that channel 1 binds, then channel 2 binds, followed by both of them unbinding. That trajectory might look like [1 1 ; 2 1 ; 2 2; 1 1]. A search with a baseline of [1 1] would find that event, and group it with all others of its kind. Setting a baseline is optional; a search can be run with one baseline, the baseline later edited, and the analysis run again to find all events matching the new baseline criteria. The baseline can be any configuration, and wildcards can be specified for one or more of the states to give the search more flexibility. Thus if the baseline is set to [Any 1] (because the researcher sees the activity of the second channel as being sufficient to define an event), then events such as [ 2 1 ; 2 2 ; 1 2 ; 1 1 ] and [1 1 ; 1 2 ; 1 3 ; 2 3 ; 2 1] will both be counted as matches. However, in any case, these various possible events are grouped together as one "class", but are also split into the various specific event types which are represented within the vague classification. So, all events matching the pattern [1 1 ; 2 1 ; 2 2; 1 1] will be grouped together, and a separate search will be made for [1 1 ; 1 2 ; 2 2; 1 1].

Setting the time step: if you want your results to be in units of “seconds”, then the number entered into this box should correspond to the time interval (in seconds) between each frame of the data imported. The analysis results will not automatically refresh, and the “Analysis→Run/Refresh Analysis” operation must be run again to update all the output variables.

Updating the filetype: if you created a KERA session, then later downloaded an updated version of the software, you should still be able to open it in the new version, but if you do, it is recommended that you run the “Update File” command under the settings tab. This is to prevent any uninitialized variables in the old version from causing errors when the new version tries to call them. If you plan to make edits to the code which might break older versions of Kera sessions, it might be a good idea to add some error handling to the updateOldFile function at the end of the Kera.m methods section.

**Appendix:**

Regular expressions (regex/regexp) search:
“Regular expressions” is a type of text search which has a much more flexible array of wildcard, quantifier, and conditional operators than the simple “NaN” and “Inf” flags used for convenience in the original KERA search style. For researchers who have a more advanced concept of the type of event they would like to search for. Some examples for a two-channel system are below:

<table>
<tr><td>Expression:</td><td>Interpretation:</td></tr>
<tr><td>[ \d  \d</td><td>The set of all configurations occurring at the beginning of a trace (the length of time “before something happens” Here, the square bracket is being used as a literal character, not a regex operator</td></tr>
<tr><td>( \d  3 ; \d  4 ;)+( \d  3)?</td><td>All events where channel 2 just alternates between state 3 and state 4 (while channel 1 stays constant at any state)</td></tr>
<tr><td>( [^1]  [^1] ;)+</td><td>All contiguous strings of configurations where neither channel is at state 1.</td></tr>
<tr><td>(( \d  [^1] ;)| ( [^1]  \d ;))+</td><td>All contiguous strings of configurations where at least one channel is not at state 1.</td></tr>
<tr><td>(?<=1  1 ;) ( [^1]  [^1] ;){3}(?= 1  1)</td><td>All events between ground states which are exactly three events long and in which neither channel is at the ground state.</td></tr>
</table>

The above examples are all a bit specific and contrived, in order to showcase the possibilities for the flexible language which regex allows the user to employ.

In order to run a regex search, the dataCell of states needs to be turned into a single string. Each cell of dataCell is acted on using the mat2str function, then the resulting strings are concatenated. A few spaces are added, and a semicolon at the end of each trace. The final string is formatted as follows:

One channel:
“[ 1 ; 2 ; … ; 3 ;][ 2 ; 1 ; … ; 1 ;]…”

Two channel:
“[ 1  1 ; 2  1 ; … ; 3  2 ;][ 2  2 ; 1  2 ; … ; 1  1 ;]…”

(The ellipses above are not part of the literal string, and are only added to show where part of each trace, and subsequent traces, have been omitted for clarity)

The number of spaces between each numerical entry is important; between channels at the same time point, there will always be two spaces; after the opening bracket at the beginning of each trace, and surrounding the semicolons which separate the time points, one space is used. If there are no results for a given regex search, checking the spaces included in your search term might be the first place to start.  Information about how regex is parsed can be found using online resources like regexr.com or the MATLAB documentation for the regexp function.  A few tips for getting started are below:

<table>
<tr><td>Expression fragment:</td><td>Interpretation:</td></tr>
<tr><td>\d</td><td>A wildcard for a state; in regex, this stands in for any digit character (0-9); if you have any traces with a state which is 10 or greater, you’ll have to use \d+ as the wildcard instead, so it covers both characters which make up two-digit numbers, but can still match single-digit characters.</td></tr>
<tr><td>[^1]<br>[^2]  etc.</td><td>An expression matching any single-character state which is not 1, not 2, etc.</td></tr>
<tr><td>(1|2)<br>(4  3|3  4)</td><td>The vertical pipe character | is used as a logical “or”; any grouping (or even a whole expression) on either side of | allows either side to be matched.  So, (1|2) matches 1 or 2, (4  3|3  4) matches either a transition from 4 to 3 or from 3 to 4</td></tr>
<tr><td>( \d  \d ;)+</td><td>Encapsulating a time-point, along with the semicolon, in a grouping parentheses allows a quantifier (like the + in this case) to be applied to the whole thing.</td></tr>
<tr><td>[^\]]+? …</td><td>A quantifier which works a bit like the “Inf” flag from the more user-friendly search interface.  This will continue to match states and transitions until either the end of the trace is hit (in which case the match will be discarded) or the expression placed after this portion (the … in the example) is matched.  So, “1 [^\]]+? 1” will match any sequence which states at state 1 and eventually returns to state 1.  For a multi-channel example, consider “1  2[^\]]+?1  2”, which works just like a “1 2; Inf Inf; 1 2” search from above.</td></tr>
<tr><td>[</td><td>The beginning of each trace.  The ‘[‘ and ‘]’ characters are special characters in the regex language, but when they are unpaired they do not need to be escaped.  You can place expressions into a parser such as regexr.com to make sure that any literal brackets you would like to include are treated as such.</td></tr>
<tr><td>]</td><td>The end of each trace</td></tr>
<tr><td>(?<=1  1 ;)…</td><td>This positive lookbehind means that only expressions which come after a [1  1] will be matched, but the [1  1] state itself will not be counted in the total time of the event.  The placement of the semicolon is important: if</td></tr>
</table>

| | |
|---|---|
|  | you don't want the [1 1] time point to be included in the final match the semicolon needs to be in the lookbehind. |
| …(?=1 1) | A lookahead: the provided configuration must happen *after* the rest of the search term, but will not be included in the final accounting of the event. The semicolon does not need to be included (but can be). |
| {2,} | A quantifier which means the previous statement must be matched two or more times. |

**Step-by-step Example:**

(?<=(2|3)  \d ; )((1  2 )|(1  1 ; 1  2 ))(; 1  \d )*(?=; (2|3))

| Before start of event: | Start of Event | | Event continues (optional) | After end of event |
|---|---|---|---|---|
| Channel 1: 2 or 3<br>Channel 2: Any | Channel 1:<br>State 1 | Channel 2<br>Either starts at state 2 or transitions there while channel 1 still at state 1 | Channel 1 still at state 1, channel 2 doesn't matter (repeat 0 - ∞ times) | End the * wildcard operator when channel 1 transitions to either 2 or 3 |

This expression was written for a two-channel model where the first channel had 3 states and the second channel had 2.  It was used to find the length of all regions during which channel 1 was at state 1 and *at some point* during that state, channel 2 was in state 2.  First, the positive lookbehind says that we want to start keeping track of the event *after* channel 1 was in one of the other states; this ensures that we aren't starting at the beginning of a trace, where the actual length of the state-1 region is unknown because we don't see the start of it, but the length of that previous state will not be included in our final accounting of the time length.  Note that the OR operator is used to say that channel 1 could be in state 2 OR 3, and the wildcard \d is used to say that we don't care about the state of channel 2 there.  Then an OR operator is used in the next group to indicate the two possible ways our scenario could occur: either channel-1-state-1 starts *while* channel 2 is at state 2, OR channel-1-state-1 happens initially while channel 2 is still at state 1, but the *following* transition sees channel 2 going to state 2.  Those are the only two ways the event can start which allow it to meet our criteria.  Then we want to continue following the event without knowing how many more transitions will occur before channel 1 stops being state 1, so we create an optional capture group that will repeat as long as channel 1 is at state 1 (so channel 2 could be fluctuating multiple times during this state) and only end it when the *next* transition sees channel 1 at state 2 or 3 (positive lookahead).

This is just one example of the versatility of the regex language in encoding a complex pattern or requirement into a simple string; full reference for regex is provided elsewhere, by more competent resources.

**Import file standards:**

The example files included with the repository in the folder called "Example input files" contain a readme and more information about the format of each file.  Examples of one-channel and two-channel datasets are included.  All input files are converted to a common format in the "dataCell" field.  This field is described in more detail inside of the Kera class methods.  There are also added instructions on how to add a custom Import function for data types that are not currently supported. By modifying the indicated lines in the openKera.m and Kera.m files, a custom import option can be added to the user interface.

Raw import specifications: the dataCell must be an N-by-Channels-by-2 cell array, where N is the number of traces (or trace pairs, etc.) and Channels is the number of channels. Each cell must contain a column vector, and the colunm vector in importedData(1,a,1) must have the same length as importedData(1,b,1) and importedData(1,a,2). This is because importedData(1,a,1) and importedData(1,b,1) are the time series of the same colocalized trace but in the different channels. The (:,:,1) entries are the "raw" data values (the pure fluorescence or FRET values) whereas the (:,:,2) entries are column vectors of integers, which correspond to the discretized traces. All values in the discretized trace cells must be integers greater than 0 in order for the "viewing" functions to work correctly, and though the analysis can probably still run if some of the states are 0, the custom search interface will not be able to access those states (you may have to dig in to the code a bit to change how the custom search works). Putting anything in the "raw" data cells is completely optional, as this only shows up when viewing the data and setting the dead time, and otherwise has no bearing on the analysis. If you have no raw data, leave the cells empty (so only the (:,:,2) cells will have any data in them).

**If MATLAB crashes:**

All is not necessarily lost. Closing the KERA window normally prompts the user for a save option; either the user selects “yes” to close the KERA window, or “no” and the window will not close. However, if the window is closing because of an internal problem or a crash, and you see this screen, you might want to click the third option, which will attempt to (before MATLAB goes completely under) save everything you had in your workspace to a single file. Later on, you might be able to get your KERA session back by opening this workspace file, finding the variable that stored your Kera, saving that variable to its own file, and then importing it as a saved session. This feature was included for the eventuality that a user’s Matlab throws some error which has the unlikely consequence of closing all windows and restarting the program; the rescued data will be sent to a file called “savingThrowError.mat” and saved wherever the last working directory was.

However, if MATLAB’s crash is so abrupt that you didn’t even see the closing prompt window (for example, if it receives a “force quit” or “end task” signal from the system), there’s not much you can do. That Kera session is lost and you’ll have to start back from whatever your last saved session was.

**Reporting bugs and contributing:**

KERA is available as a public GitHub repository, which means that anyone with a GitHub account can become a contributor, report bugs, and suggest changes. You can create your own copy of KERA and edit it to your liking, and request that changes be pulled into the Main branch of the original, to benefit others who download it. Reporting bugs to the “issues” page might prompt a fix, and suggesting new features there would also be welcomed. If you plan to make edits to the code which might break older versions of Kera sessions, it might be a good idea to add some error handling to the updateOldFile function at the end of the Kera.m methods section; this way, if someone sends you their KERA saved session, you can quickly update it to work with your version.

**SUPPLEMENTARY NOTE 2**
### Normalization Scripts

Most of the popular discretization programs are written with an assumption of the FRET experiments. Fluorescence data in individual channels, which are not from FRET experiments need to be modified to be used in such programs.

The main purpose of the normalization scripts is to read in column data from multiple files (in formats such as .txt, .csv, and .dat) and normalize each trajectory to occupy values between 0 and 1. In this way, fluorescence level data may be compared at a common "scale" or input to programs such as ebFRET which require normalized data.

The main function is normalizeTrajectorySet.m, which is located in the main folder of the repository. It contains calls to functions inside the "functions" directory, so the working directly should be set to the sourceCode folder before running (otherwise the user will be prompted to switch directories). The user is first prompted for the number of channels. As described in the KERA documentation, channels are independent signals. In a dual-illumination experiment observing Cy3 and Cy5 simultaneously, and colocalizing the signals, there would be two channels. In a study with only one kind of fluorophore, there is likely only one channel. This was the case for the RPA study (8). Then, the user is prompted to select a directory which contains data files. The program assumes that all files of a certain type (specified by the user as .txt, .csv, or .dat) in that directory contain columns of data which should be imported. One column from each file is imported, according to the column the user specifies. For example, many .dat files contain two columns: a time series and an intensity trajectory. In that case, the user would specify that column 2 contains the desired data. If there is more than one channel, the user is prompted to go through these steps for each channel. The goal is to keep associated (colocalized) trajectories sorted in the same order; assuming the filenames are consistent, this is done automatically by the dir function upon input, especially if the colocalized signals are represented by different columns in the same files. If your data is in some other form, rearranging it into the format found in the intensity and fileNames variables from the sample data folder (normalizationSampleData.zip). By loading it in and setting "importOldSession" to 1, the default importation steps may be skipped (you may have to manually assign some other variables, like channels).

Once the data is imported, the user is prompted to view the traces. This is a good idea, unless the viewing/trimming process has already been completed for this data set and you are working from old imported data. Upon viewing, the user is guided through each trajectory in the set sequentially, viewing all channels (if there is more than one) at once, but only editing them one at a time. So, trajectory 1 channel 1 is seen first, followed by trajectory 1 channel 2, and so on. For each trajectory, the user can perform four actions:

- Discard the trace: this removes it from analysis, and also removes the associated trajectories in all other channels (if it is a multi-channel data set).
- Trim trace: clicking this button allows the user to select two points (the start and end) of the portion of the trajectory they would like to keep. In a multi-channel dataset, the corresponding trajectories from all channels will be trimmed to this

length. The appearance of the data does not change to reflect the trimming. Clicking this button again on the same trace overwrites the previous trim region.

- Select low: select a region by clicking twice (start and end points) which represents a continuous region of the “low” state of the trajectory. This is the state which will be set at the baseline when the trajectories are aligned and normalized. Clicking this button again on the same trace overwrites the previous low region selection.
- Select high: select a region in the same way, but which should be normalized as a “high” value (aligned to 1 or near 1). Clicking this button again on the same trace overwrites the previous high region selection.
- Go back: go to the previous trajectory

If no trimming is done, the full length of the trajectory will be included in the analysis. A feature reminds the user to trim, select low, and select high on every single trace, but this can be suppressed by clicking the appropriate option on the dialogue box. It is still recommended that at least every trace be assigned a low and a high region; this is because automatic normalization is highly data-dependent and the assumptions made by that algorithm may not be appropriate for your data.

When all traces have been viewed, the user is taken back to the main code, and a full export package is saved; an example of this is in the zipped file normalizationSampleData.zip in the sample data folder. It holds the original imported data as well as the edits made during the viewing session. Saving at this point means the user can always come back and make additional edits (running the code with importOldSession set to 1 after opening the .mat saved variables), either in the view window or in the normalization.

If all traces have had a low or high region selected, the normalization proceeds entirely automatically from that point, and the last step is to select the export formats and export locations. The original filenames are appended with flags specifying the filetype and saved in the locations indicated by the user. hFRET requires that all data within a set be the same length; if the trajectories have been trimmed to non-uniform lengths, the normalization script will ask the user for a padding value which will be appended to all trajectories until they are the same length. By setting this to a value like -.5, the user can assure that hFRET will assign the padded values to their own state (and only the padded values to that state) though this does mean that the hFRET parameters must be set to fit an additional state than would normally be part of the model. However, these details are only relevant to post-processing and dependent on the discretization program the user is planning to import the normalized traces to.
